# Highly potent novel multi-armoured IL13Ra2 CAR-T subverts the immunosuppressive microenvironment of Glioblastoma

**DOI:** 10.1101/2025.01.10.632392

**Authors:** Maurizio Mangolini, Saket Srivastava, Emily Souster, Yajing Yang, Huimin Wang, Rajesh Karattil, Liam Schultz, Biao Ma, Diana Pombal, Matthew Greenig, Aubin Ramon, Pietro Sormanni, Shaun Cordoba, Shimobi Onuoha

## Abstract

Glioblastoma remains one of the most challenging and lethal brain cancers, with limited treatment options. While CAR-T cells have shown promise in some patients, sustaining T-cell activity and overcoming the immunosuppressive tumour microenvironment remain significant hurdles. Here, we present an armoured CAR-T cell design to address these challenges and enhance persistence in GBM tumours. We developed a highly specific humanised single-domain antibody targeting IL13Rα2 and included it alongside four additional modular elements in a single retroviral vector for CAR-T generation. Our results demonstrate that this single-cassette CAR-T cell design possesses high resilience against TGF-β-mediated immunosuppression, enhanced tumour-killing capacity through IL-12 secretion while maintaining a favourable safety profile, extended persistence in the host, and an additional layer of safety control through the incorporation of a suicide switch. Importantly, despite its complexity, the construct can still be manufactured efficiently. These advancements represent a significant step forward in addressing key challenges associated with CAR-T cell therapy in solid tumours.

## INTRODUCTION

Glioblastoma (GBM) is one of the most aggressive and lethal forms of brain cancer, with a dismal prognosis despite current standard treatments and the implementation of multimodal treatment strategies. Unfortunately, at present day, the prognosis for glioblastoma remains poor with a median survival of less than 15 months from initial diagnosis, a 2-year survival rate of approximately 26-33% and a 5-year survival rate estimated at just 6.8%^1,2^.

In recent years, chimeric antigen receptor (CAR) T cell therapy has emerged as a promising immunotherapeutic approach for treating GBM^3^. Among the various targets explored, interleukin-13 receptor alpha 2 (IL13Rα2) has garnered significant attention as a potential CAR-T cell target for this disease indication. IL13Rα2 is a monomeric high-affinity receptor for interleukin-13 (IL-13) that is overexpressed in most GBM tumours, with studies reporting its presence in 60-75% of cases. Importantly, IL13Rα2 expression is minimal in normal brain tissue, making it an attractive tumour-specific target^4,5,6^. The receptor’s overexpression is associated with the mesenchymal GBM subtype and correlates with poor patient survival, further underscoring its relevance as a therapeutic target^7^.

CAR-T cells engineered to target IL13Rα2 have shown promising results in preclinical studies triggering a potent anti-tumour immune response. Furthermore, recent clinical trials evaluating IL13Rα2-targeted CAR-T cells in GBM patients have shown encouraging results, with evidence of tumour regression and improved survival in some cases^8,9,10,11^. However, challenges such as limited T-cell persistence and exposure to the immunosuppressive tumor microenvironment (TME) have been observed. One way to overcome the challenges faced by CAR-T cell therapy in treating solid tumours like GBM is to develop “armoured” CAR-T cells with enhanced capabilities.

Similarly to most solid tumours, GBM employs various immune-evasive strategies within their tumour microenvironment (TME) that result in treatment resistance. These mechanisms pose substantial challenges for CAR-T cells, hindering their ability to mount effective antitumor responses and achieve the robust expansion necessary for significant tumour reduction. Among these, the Transforming Growth Factor beta (TGF-β1/TGF-βR2) signalling pathway is particularly significant in promoting T cell dysfunction and limiting their proliferation^12,13,14^. Enhancing CAR-T cells with the ability to produce anti-tumour cytokines is also a promising strategy to overcome the challenges posed by the hostile TME. Interleukin-12 (IL-12) stands out among these cytokines for its remarkable anti-tumour capacity. However, the clinical application of IL-12 has faced significant hurdles due to its potent systemic toxicity^15,16^. Indeed, multiple experimental models and clinical trials have revealed toxicity profiles associated with systemic IL-12 exposure, which has hampered its progression in therapeutic development^17^.

Here we show a novel armoured CAR-T cell targeting IL13Rα2, which addresses several key challenges associated with CAR-T cell treatment of GBM. Our approach combines a newly developed, highly specific, humanized single-domain antibody (VHH) against IL13Rα2 with a fourth-generation CAR design that incorporates multiple functional enhancements within a single retroviral vector. This advanced CAR construct includes: -a TGF-β receptor-blocking antibody; - a constitutively active form of the GM-CSF intracellular receptor region; -an engineered single-chain IL-12 sequence; - a modified version of HER2 as a suicide switch for controlled cell elimination. We demonstrate that primary T cells expressing this construct show superior performance compared to conventional CAR-T cells, exhibiting enhanced tumour cell killing, improved proliferation, and prolonged persistence. The blockade of TGF-β signalling significantly enhances the engineered T cells’ resistance to immunosuppression, while the attenuated IL-12 activates bystander NK and T cells without observable toxicity. Furthermore, the incorporation of a suicide gene, activated by the FDA-approved drug Trastuzumab emtansine (T-DM1), provides a crucial safety mechanism for the controlled elimination of the engineered cells. *In vitro* and *in vivo* data confirmed the efficacy, survival, and safety profile of these armoured CAR-T cells, even at low doses. Notably, despite the complexity of the construct, we demonstrate manufacturing feasibility in a Good Manufacturing Practices (GMP) - compliant setting. Altogether, our findings offer a promising new avenue for the treatment of GBM and potentially other IL13Rα2-expressing malignancies representing a significant step forward in the field of CAR-T cell therapy for solid tumours.

## RESULTS

### *In vitro* screening identifies lead IL13Ra2 VHH for CAR-T cells

The targeting domain of CARs is their most critical component for recognizing and interacting with the desired target antigen. To develop a new CAR targeting IL13Rα2, we created a specific nanobody, due to their advantages in terms of low immunogenicity and small size^18^. The discovery of the IL13Rα2-specific nanobody was achieved through a combination of immunization, phage display, and deep repertoire mining, followed by humanization and functional screening (Figure 1A). In brief, after immunizing alpacas, we generated a phage display library enriched against IL13Rα2, using IL13Rα1 as a deselection target to ensure high specificity for IL13Rα2 while avoiding cross-reactivity with IL13Rα1.

**Figure 1:**
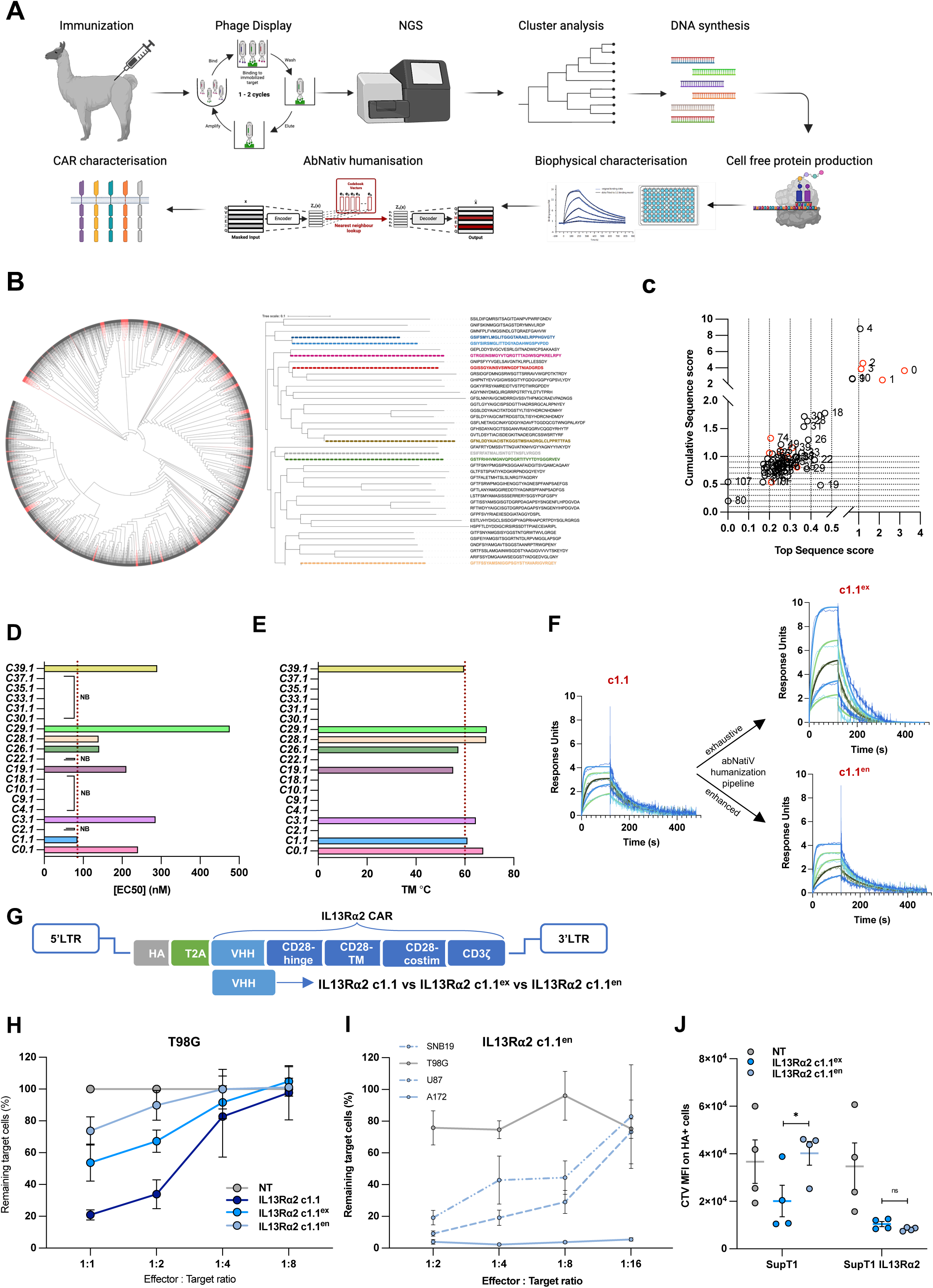
I*n vitro* screening identifies lead IL13R⍺2 VHH for CAR-T cells. **A.** Graphical scheme of the experimental procedures to identify a new IL13Ra2 VHH for CAR-T cells created with BioRender.com. **B**. Phylogenetic tree generated using iTol, illustrating the complete library diversity. Clusters identified during analysis are highlighted in red. Clusters were determined by concatenating the three CDRs and analysing their diversity using a chemical space distance matrix. The sequences of the top clusters were further analysed in a unique tree to explore the distances between the combined CDRs of the highest-represented clones within the diverse clusters. **C**. Analysis of potential library bias during panning, represented as a plot of the top sequence score against the cumulative sequence score, highlighting the enrichment dynamics and selection preferences. Bar plots of the binding values (**D**) and the apparent melting temperatures (**e**) of the selected clones produced using cell-free based expression system. Values are representative measurements from three independent replicates. NB: non-binding. **F**. Binding sensorgrams of respective representative measurements showing the binding of c1.1, c.1.1en and c1.1ex to human IL13Rα2 by SPR. IL13Rα2 c1.1ex and IL13Rα2 c1.1en are based on the selected c1.1 clone and humanised using the exhaustive (ex) or enhanced (en) abNatiV pipeline. **G.** Schematic representation of CAR-T vectors targeting IL13Rα2. The vectors comprise an N-terminal HA-tag serving as a transduction marker, followed by the anti-IL13Rα2 VHH of interest. The VHH is linked to the transmembrane and intracellular regions of the CD28 signalling domain, which is then connected to the CD3ζ signalling domain. LTR, long terminal repeat. **H,I.** Different glioma cell lines were co-cultured with different ratios of CAR-T cells for 72 hours. Viable target cells were quantified by flow cytometry using CD2 and CD3 expression as exclusion markers. The percentage of live target cells was calculated by normalizing to the NT group. **J**. Antigen-specific proliferation was determined by co-culturing CTV-labeled T cells with SupT1-IL13Rα2 or SupT1 cells for 96 hours. The T cells were either untransduced (NT) or expressing the IL13Rα2 c1.1en or IL13Rα2 c1.1ex CAR vector. *P < 0.05, **P < 0.01, ***P < 0.001, ****P < 0.0001 and not significant (ns) P > 0.05.

Next-generation sequencing (NGS) was then performed on the enriched library, and cluster analysis identified potential binders. The analysis focused on extracting complementarity-determining regions (CDRs) 1, 2, and 3, which were computationally combined to generate artificial sequences for similarity clustering, with the idea that VHHs with similar CDRs should bind the same epitope. Homology groups were formed using a distance matrix that prioritized chemical differences and similarities between amino acids, rather than simpler sequence-identity (Supplementary Figure S1A). The highest frequency sequence from each group was selected for further characterisation. This approach effectively sampled the diversity of the screened library and reduced the risk of clonal dominance.

Parallel to NGS analysis, traditional panning and clone selection by colony picking were also performed, yielding complementary insights. A total of 50 clusters were initially identified, with 20 clusters progressing to production and subsequent functional evaluation (Figure 1B,C). Binding specificity assays identified eight molecules that bound IL13Rα2 without cross-reactivity to IL13Rα1. However, despite the extensive diversity of the library, repeated rounds of panning narrowed the functional binders to two primary sequences. This observation suggests a potential bias in phage display selection that was not mirrored in the broader diversity identified through NGS.

Half maximal effective concentration (EC_50_) and thermal stability (T_m_) analysis revealed that the majority of the selected binders exhibited favourable melting temperatures, indicative of robust biophysical properties in addition to specific binding (Figures 1D,E). Based on these combined biophysical and functional analyses, clone C1.1 from cluster 1 emerged as the lead candidate for incorporation into the IL13Rα2 CAR, designated IL13Ra2 c1.1.

We then performed humanisation of the Clone c1.1 using the AbNatiV VHH humanisation pipeline, as reported in Ramon et al. ^19^. This AI-based approach samples humanising mutations ensuring retention of binding affinity and favourable VHH biophysical properties (Figure 1F). The humanized variants c1.1^en^ and c1.1^ex^, respectively from the enhanced and exhaustive AbNatiV humanisation pipelines^19^ showed equivalent stability and specificity compared to the wild-type counterpart. To further assess the specificity of these VHHs for IL13Rα2 and minimize the risk of off-target binding effects, we conducted an extensive cross-reactivity study. This study evaluated the binding of the VHHs against approximately 6,000 native human membrane proteins. Remarkably, all three clones demonstrated minimal cross-reactivity with other proteins. Among them, c1.1^en^ exhibited the strongest specific binding to IL13Rα2 (Supplementary Figure S1B).

Clones c1.1, c1.1^en^ and c1.1^ex^ were then inserted into a retroviral vector encoding a CAR construct with CD28 and CD3ζ intracellular signalling domains (Figure 1G). These were tested against the T98G glioma cell line (IL13Rα2-negative) and SUPT1 cells engineered to express IL13Rα1 to assess off-target and cross-reactivity. Notably, the humanized variant c1.1^en^ exhibited minimal off-target activity against IL13Rα1, corroborating its specificity and reduced background activity or “tonic signalling” compared to both the wild-type llama-derived CAR and the exhaustive-humanized c1.1^ex^ CAR while retaining high cytolytic activity against IL13Rα2-expressing glioma cell lines and SUPT1-IL13Rα2 cells (Figure 1H,I and Supplementary Figure S1C-G). This reduction in background activity is likely due to improved biophysical properties, such as increased solubility and thermal stability, engineered during the humanization process. Further validation was conducted using CellTrace Violet (CTV) proliferation assays on IL13Rα2-positive and IL13Rα2-negative SupT1 cells (Figure 1J). These results highlight the advantages of the engineered humanized clone C1.1^en^, demonstrating its suitability for clinical development and its potential to mitigate adverse effects associated with CAR tonic signalling.

### dsFvTBRII/CCR overcomes the inhibitory effects of TGF-b and induces CAR-T cell persistence

TME-induced T cell inhibition and reduced persistence are among the major factors limiting the efficacy of CAR-T cells in solid tumours^20^. TGF-β, abundantly secreted by tumor and stromal cells in glioblastoma, potently suppresses CAR-T proliferation, drives exhaustion, and fosters an immunosuppressive microenvironment, thereby limiting therapeutic efficacy^21,22,23^. To overcome these limitations, we designed a fusion protein containing two key elements: an extracellular region consisting of a blocking antibody for the TGF-β receptor II (TBRII) and the transmembrane and truncated intracellular domains of the granulocyte-macrophage colony-stimulating factor (GM-CSF) receptor as a chimeric costimulatory receptor (CCR) (Figure 2A). This innovative architecture leverages the disulfide bond of the TBRII antibody region to achieve constitutive downstream activity of the receptor. We first validated the blocking functionality of the disulfide-stabilized single-chain variable fragment (dsFv) antibody derived from the original Fab sequence. Using a reporter cell line that quantifies TGF-β response through secreted embryonic alkaline phosphatase (SEAP) production, we demonstrated that the dsFv antibody retained the TGF-β blocking capacity of the original Fab version (Figure 2B). Subsequently, we evaluated the function of the intracellular component. We generated two single cassette constructs expressing the dsFvTBRII region fused with different orientations of the GM-CSF receptor chains: alpha-beta and beta-alpha. The overexpression of these constructs in Jurkat cells revealed that the beta-alpha orientation was more potent in activating STAT5 (Figure 2C). This finding was further corroborated using GM-CSF reporter cell lines (Supplementary Figure 2A). Moreover, we confirmed that the final chimeric receptor design (dsFvTBRII/CCR) maintained its TGF-β blocking functionality when tested using TGF-β reporter cells (Figure 2D). To assess whether the dsFvTBRII/CCR module would enhance CAR-T cell resistance to TGF-β and improve their persistence, we integrated this construct into the IL13Rα2 c1.1^en^ CAR design, separating the two components with a T2A self-cleaving peptide. Comparative analysis revealed that CAR-T cells incorporating the dsFvTBRII/CCR module exhibited reduced susceptibility to TGF-β pathway activation, measured by decreased SMAD2/3 phosphorylation (Figure 2E and Supplementary Figure S2B). Furthermore, in cytokine-deprived *in vitro* culture conditions (Figure 2F), control cells incurred complete cell death within 10 days, whereas cells overexpressing the CCR maintained viability for up to 40 days (Figure 2G). To determine whether the extended viability observed in CAR-T cells expressing the CCR module was accompanied by enhanced proliferative capacity, we conducted a dye dilution assay to track cell division over 7 days. Our results demonstrated that CAR-T cells incorporating the CCR module underwent multiple rounds of proliferation, significantly outperforming the control cells (Figure 2H). This finding was further corroborated by the rapid increase in the proportion of CCR-expressing CAR-T cells within the total cell population over time (Figure 2I). The long-term cytotoxic function and proliferative capacity of CAR-T cells were assessed in vitro over 7 weeks, with additional glioma tumor target cells added weekly to the co-culture. The CCR-expressing CAR-T cells maintained full cytolytic activity for at least 5 weeks, compared to only 3 weeks for non-armored CAR-T cells (Supplementary Figure S2C). To further validate the efficacy of our dsFvTBRII/CCR module, we employed the well-characterized CD19 CAR-T cell model as an additional experimental system. *In vitro* experiments demonstrated that overexpression of the dsFvTBRII/CCR construct in CD19 CAR-T cells conferred partial protection against the cytotoxic inhibitory effects of TGF-β (Supplementary Figure S2D). Moreover, we observed that cells incorporating this additional module displayed extended longevity compared to those expressing only the CAR construct, suggesting that the ability of the dsFvTBRII/CCR to enhance CAR-T cells’ persistence is independent of the upstream CAR architecture (Supplementary Figure S2E,F). Collectively, these results indicate that the incorporation of the dsFvTBRII/CCR module confers enhanced resistance to TGF-β-mediated immunosuppression and significantly extends the survival and the proliferative potential of CAR-T cells in the absence of exogenous cytokine support, potentially overcoming some of the challenges of the tumour microenvironment.

**Figure 2:**
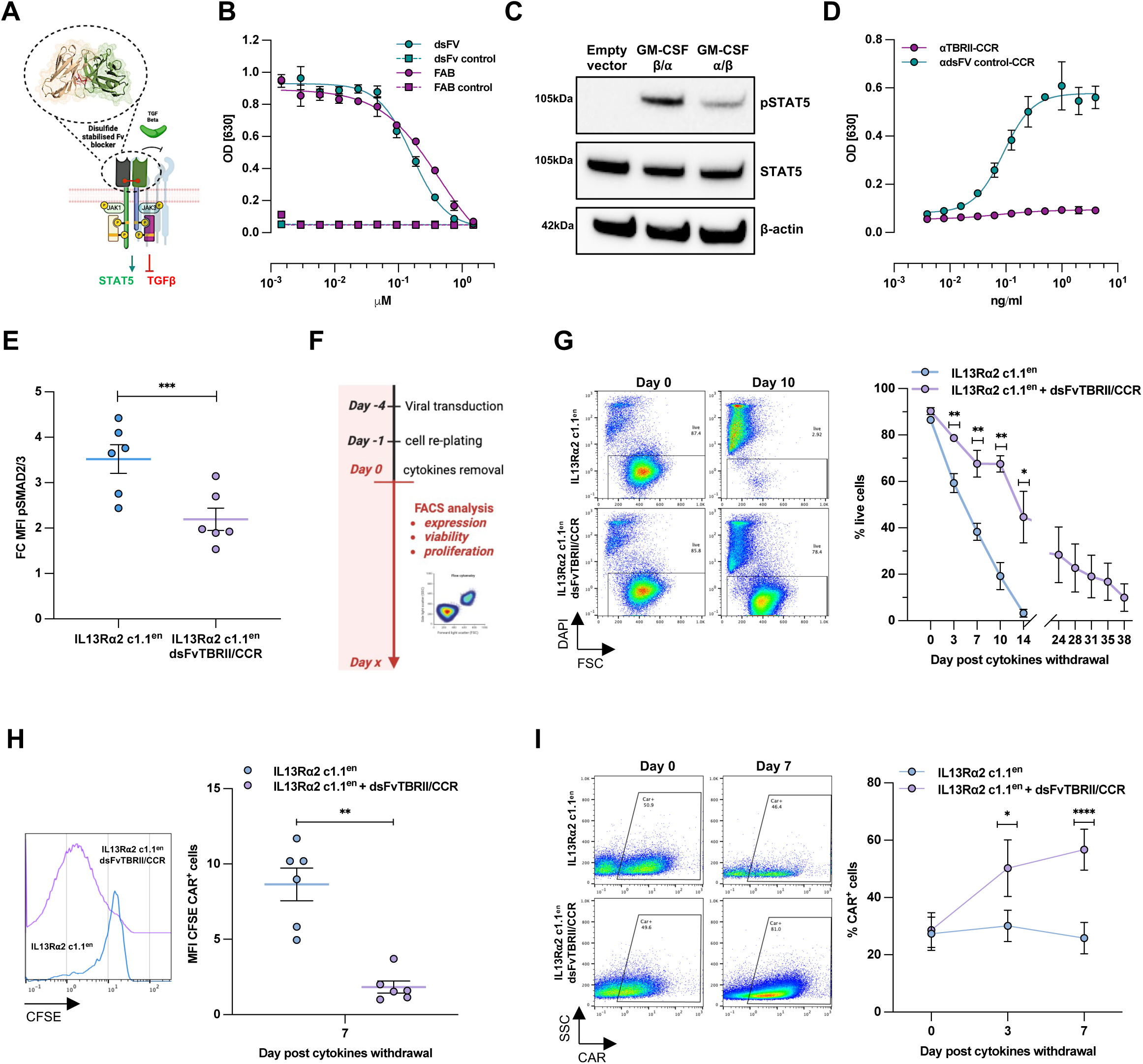
dsFvTBRII/CCR overcomes the inhibitory effects of TGF-b and induces CAR-T cell persistence. **A.** Graphical scheme of the dsFvTBRII/CCR construct. **B.** Quantification of the SEAP secreted by HEK-Blue™ TGF-β cultured in the presence of TGF-b and the indicated level of the different TBRII blocking antibodies. **C.** Analysis of phospho-STAT5 and total STAT5 levels in Jurkat cells expressing the dsFvTBRII region fused with different orientations of the GM-CSF receptor chains: alpha-beta and beta-alpha. **D.** Quantification of the SEAP secreted by HEK-Blue™ TGF-β cultured in the presence of TGF-b and the indicated level of dsFvTBRII/CCR or a non-targeting/CCR control antibody. **E.** Quantification of intracellular phospho-SMAD2/3 levels by flow cytometry in T cells expressing IL13Ra2 c1.1en CAR or IL13Ra2 c1.1en CAR+dsFvTBRII/CCR exposed to 10ng/ml TGF-β for 30’. Cells were grown in serum-free media overnight before TGF-β stimulation. **F.** Schematic representation of the experimental setup. **G.** Percentage of live IL13Ra2 c1.1en CAR or IL13Ra2 c1.1en CAR+dsFvTBRII/CCR cells following cytokine withdrawal (n=6). Live cells were identified by DAPI exclusion by flow cytometry. **H.** Measurement of transduced T-cell proliferation by CFSE dilution 7 days post cytokine withdrawal (n=6). Cells were labelled with CFSE at day 0. **i.** Analysis of CAR-positive cell enrichment over time following cytokine withdrawal measured by flow cytometry (n=6). A representative flow cytometry plot from an individual T cell donor is presented for **G**,**H**,**I**. Cohorts are shown as mean ± SEM. Statistical analysis was done by paired t-tests. *P < 0.05, **P < 0.01, ***P < 0.001, ****P < 0.0001 and not significant (ns) P > 0.05. **A**,**F** were created with BioRender.com

### Development of a scIL12 with an improved safety profile

IL-12 is a potent pro-inflammatory cytokine that is crucial in bridging innate and adaptive immunity. It has several beneficial effects in the context of cancer immunotherapy, such as enhancing T cell proliferation and cytotoxicity, activating and recruiting NK cells and NKT cells and stimulating interferon-gamma (IFNγ) production^24^. However, its potent anti-tumour effects are concomitant with severe toxicities in clinical trials, limiting its therapeutic application^17^. To mitigate potential toxicity, we compared different single-chain IL-12 (scIL12) constructs. We focused on two primary configurations that have previously shown different biological activities^25^: One comprising the p40 subunit followed by the p35 subunit (scIL12^wt^), and another with the reverse arrangement p35 followed by p40 (scIL12^inv^). Additionally, we examined the canonical G_4_S linker using different amino acid length compositions connecting these subunits. This aspect was motivated by the understanding that linker length can significantly influence protein flexibility^26^, potentially affecting the biological activity and safety profile of the scIL12 forms. To evaluate the functional differences among the various scIL12s, we assessed their capacity to activate IL-12 signalling using a commercially available IL-12 reporter cell line stimulated with scIL12 purified proteins. This system allows quantifying the activation of IL-12 signalling, measuring the production of SEAP following IL-12 binding and the subsequent activation of STAT4 (Figure 3A). The results showed that the scIL12 proteins with the p35-linker-p40 subunit configuration exhibited reduced activity compared to the p40-linker-p35 arrangement. Furthermore, shortening the linker length from 15 to 7 amino acids in the p35-linker-p40 construct resulted in an additional decrease in functional activity (Figure 3B). Notably, this difference was evident only in the p35-p40 orientation. Further investigation of the linker length in the inverted orientation revealed no correlation between shorter linker lengths and signal downregulation. Indeed, the maximum reduction in signalling was achieved using a 7-amino acid linker, suggesting that this specific length optimally decreases the biological activity of the inverted IL-12 construct (Supplementary Figure S3A).

**Figure 3:**
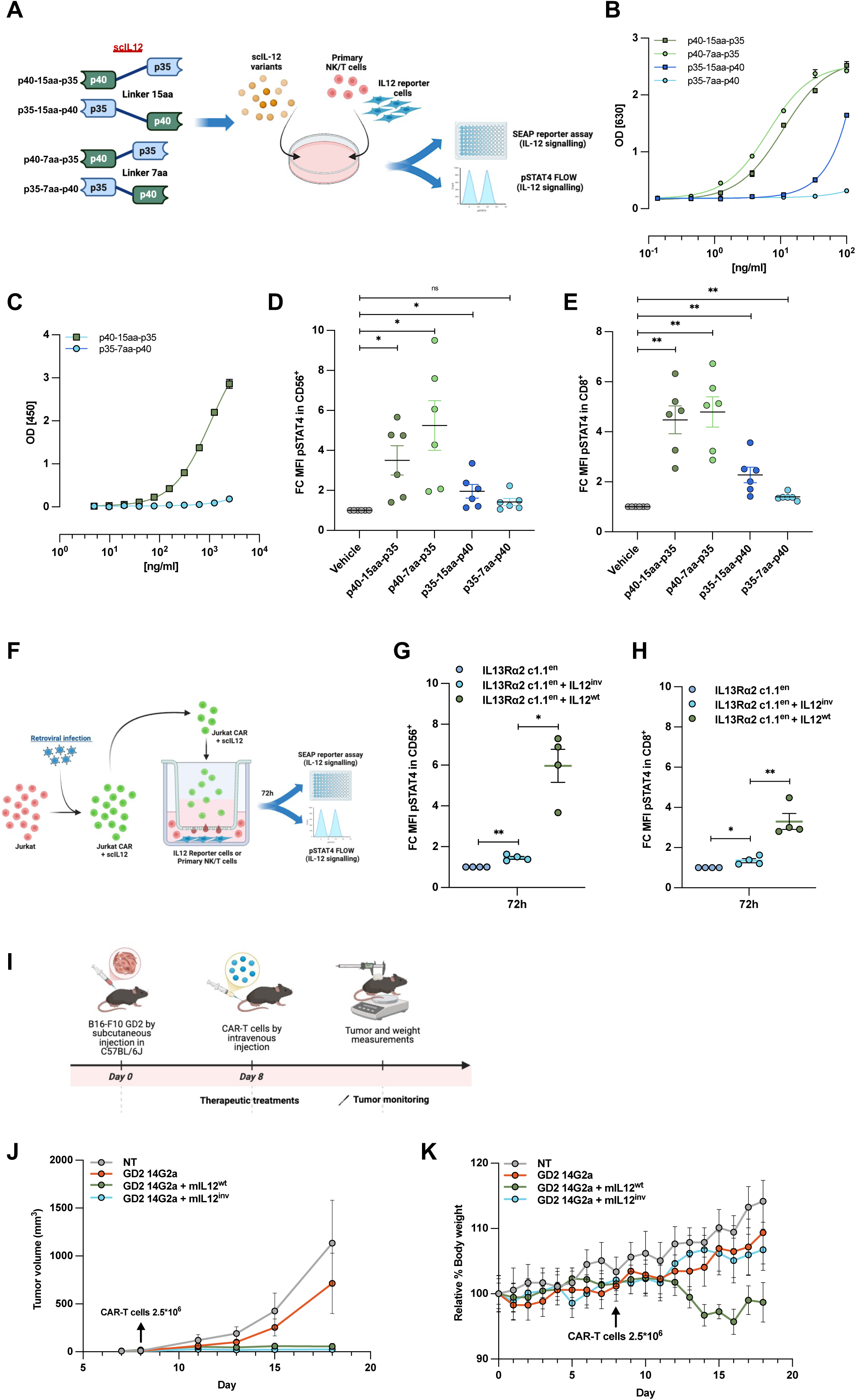
IL12inv demonstrates enhanced safety while preserving its potent anti-cancer effects. Schematic representation of the protein used and the experimental layout. **b.** Quantification of the SEAP secreted by HEK-Blue™ IL12 cultured in the presence of scIL12 purified recombinant proteins at the indicated concentration for 72h. **C.** Affinity of the designed p40-15aa-p35 and p35-7aa-p40 scIL12s to the immobilized IL12β1 evaluated by ELISA assay. scIL12s were detected using an ⍺-HIS-HRP conjugated antibody. **D,E.** CD56+ and CD8+ cell assay demonstrating signalling capacity of scIL-12 variants using purified recombinant protein at 100ng/ml for 30 minutes. Cells were harvested, fixed and permeabilised to detect pSTAT4 status by flow cytometry. T cells were previously activated with ⍺CD3/CD28 in the presence of IL-2. **F.** Schematic representation of the experimental layout. **G,H.** pSTAT4 level in CD56+ and CD8+ cells following co-culture with Jurkat cells expressing IL13Rα2 c1.1en, IL13Rα2 c1.1en+IL12inv or IL13Rα2 c1.1en+IL12inv CAR for 72h. Cells were co-cultured using a 0.4µm cell-permeable culture support. **i.** Schematic representation of the *in vivo* experimental layout created with BioRender.com. Tumor volume (**J**) and body weight (**K**) measurements of mice subcutaneously injected with 1×105 GD2-expressing B16.F10 cells, which were allowed to engraft for 8 days before a 2.5×106 dose of non-transduced T or CAR-T cells was intravenously injected via the tail vein. Cohorts are shown as mean ± SEM. Statistical analysis was done by paired t-tests. *P < 0.05, **P < 0.01, ***P < 0.001, ****P < 0.0001 and not significant (ns) P > 0.05. **a**,**i** were created with BioRender.com

Activation of the IL-12 signalling is mediated by initial ligation of the p40 subunit to the IL-12Rβ1 and subsequent heterodimerisation with IL-12Rβ2 mediated by the p35 subunit^27,28^. To investigate the impact of linker length and subunit order on the structural integrity and receptor binding affinity of our engineered scIL12 variants, we conducted an ELISA-based binding assay. Our results revealed that the orientation of the IL-12 subunits significantly influenced receptor binding. Specifically, scIL12 constructs with the p35-p40 orientation demonstrated reduced binding affinity to IL-12Rβ1 compared to those with the p40-p35 orientation (Figure 3C). Interestingly, we observed no significant differences in binding affinity to IL-12Rβ2 across the various scIL12 orientations (Supplementary Figure S3B). This indicates that the order and spatial arrangement of the subunits play a crucial role in maintaining the proper conformation for receptor interaction. To further evaluate the biological activity of our engineered scIL12, we tested their efficacy on T cells and NK cells. The results showed that STAT4 phosphorylation (pSTAT4) levels varied in a specific manner, influenced by the length of the linker connecting the IL-12 subunits and their orientation (Figure 3D,E). To validate the functionality of scIL12 in the context of CAR-T cells, we incorporated scIL12^wt^ (p40-15aa-p35) or scIL12^inv^ (p35-7aa-p40) in the IL13Rα2 c1.1^en^ CAR vector. Engineered cells were co-cultured using cell-permeable supports with primary isolated T and NK cells for 72 hours to assess IL-12-mediated activation through intracellular staining of pSTAT4 and measurement of IFNg production (Figure 3F). Our results demonstrated that while the IL13Rα2 c1.1^en^ + scIL12^inv^ construct maintained the capacity to activate both T and NK cells, its potency was notably reduced compared to the scIL12^wt^ cells (Figure 3G,H). The enhanced safety profile was particularly evident in the reduced production of IFNγ, the cytokine primarily responsible for IL-12 toxicity, by NK cells following IL-12 stimulation (Supplementary Figure S3C). Some studies demonstrate antigen-dependent regulation of IL-12 secretion in armored CAR-T cells^29^. To determine if this occurs with our specific construct, we evaluated scIL-12 expression in both wild-type and inverted configurations upon antigen stimulation. Results revealed a trend toward increased IL-12 secretion, potentially due to CAR-T activation driving enhanced proliferation and thus a greater number of IL-12-producing cells (Supplementary Figure S3D). To assess both the anti-tumour efficacy and potential toxicity of our engineered scIL12 *in vivo*, we utilized an aggressive mouse melanoma model using B16.F10 cells transduced to express GD2. We established tumours in syngeneic mice by injecting B16.F10-GD2 cells and allowing them to engraft for 8 days. Subsequently, we administered a single intravenous dose of 2.5 × 10^6^ GD2, GD2-mIL12^WT^, or GD2-mIL12^inv^ CAR-T cells (Figure 3I). Our results demonstrated that while GD2 CAR-T cells alone provided limited tumour control, both IL-12-expressing CAR-T cell groups showed remarkable anti-tumour efficacy, with tumours remaining nearly undetectable (Figure 3J and Supplementary Figure S3E). Notably, we observed a complete contrast in toxicity profiles between the two IL-12 variants. Mice receiving GD2-mIL12^wt^ CAR-T cells experienced significant weight loss, while mice treated with GD2-mIL12^inv^ CAR-T cells maintained stable body weight, suggesting a substantially improved safety profile of IL12^inv^ (Figure 3K).

In conclusion, our findings demonstrate that the engineered scIL12 with inverted subunit orientation (p35-p40) and shortened linker length maintains a significant level of biological activity while substantially reducing the systemic toxicity commonly associated with IL-12 therapy.

### HER2 as a safety switch and CAR-T detection marker

To address potential safety concerns associated with IL-12 secretion, induced enhanced proliferation and prolonged persistence of our CAR-T cells, we incorporated an elimination switch into our construct. This safety mechanism is designed to provide a means of controlling the engineered T cells *in vivo* if required^30^. For this purpose, we selected HER2 as the protein of interest due to its compatibility with an existing FDA-approved targeted therapy. Trastuzumab emtansine (T-DM1), an antibody-drug conjugate targeting HER2, has demonstrated high efficacy and is generally well-tolerated in clinical settings^31^. In our efforts to optimise the HER2-based suicide switch module, we explored various modifications to enhance its efficacy while reducing its overall length in our construct (Figure 4A). Firstly, we replaced the native HER2 transmembrane domain with that of CD5, a protein known to facilitate rapid receptor internalisation in T cells^32^. Our results showed that cells expressing the HER2-CD5 transmembrane construct exhibited significantly increased sensitivity to T-DM1 compared to those expressing the original HER2 construct. We further optimized the suicide switch by focusing on its extracellular region. We generated a truncated version of the HER2 extracellular domain, retaining only the specific region containing the docking site for T-DM1 (DHER2^CD5^). This modification was designed to minimize the size of the construct while preserving its essential function as a safety switch. Our results demonstrated that this truncation did not compromise the construct’s sensitivity to T-DM1. Remarkably, we observed an increase in sensitivity to the antibody-drug conjugate compared to the full-length extracellular domain (Figure 4B). To further validate our findings, we extended our investigation to primary T cells, comparing the efficacy of T-DM1-mediated elimination between cells expressing the IL13Rα2 c1.1^en^-DHER2^CD5^ CAR construct and those expressing the CAR alone. T cells engineered with the IL13Rα2 c1.1^en^-DHER2^CD5^ CAR construct exhibited significant susceptibility to T-DM1-induced elimination, whereas those expressing only the CAR remained largely unaffected (Figure 4C,D). We also observed a notable decrease in CAR expression intensity, suggesting that T-DM1 preferentially affects cells with higher CAR expression levels. This observation indicates that the reduction of highly expressing cells, which are likely responsible for increased toxicity, occurs first (Figure 4E). These findings support the translational potential of our optimized suicide switch strategy for clinical applications.

**Figure 4:**
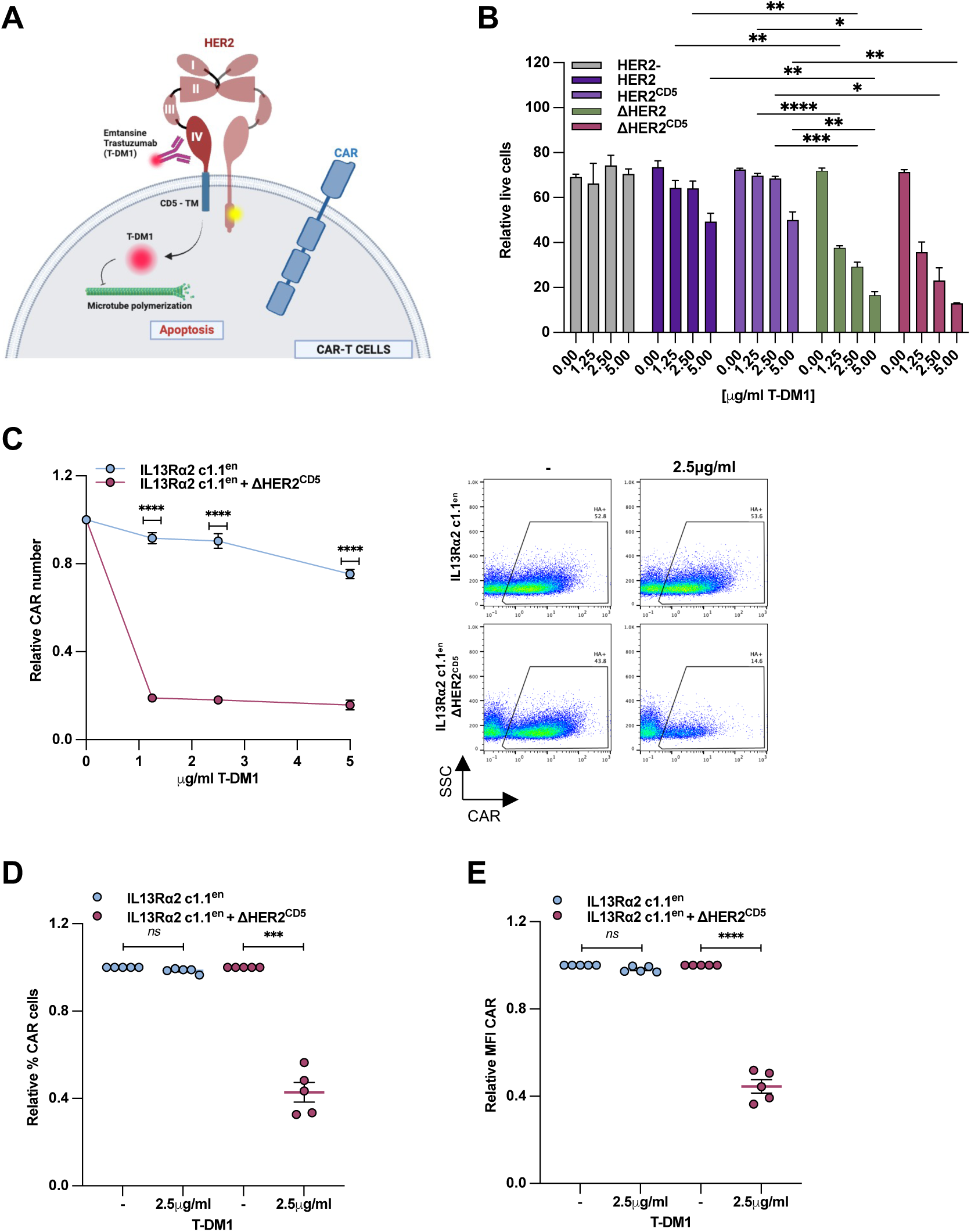
ΔHER2CD5 enhances the safety of CAR T cells as safety switch. **A.** Schematic representation of the HER2 mode of action as suicide switch created with BioRender.com. **B.** Relative percentage of live (DAPI− Annexin-V−) Jurkat cells overexpressing full length HER2, HER2 with CD5 transmembrane or truncated HER2 following T-DM1 exposure at the indicated concentrations for 72h. Results are expressed as a representative of three independent experiments. **C.** IL13Rα2 c1.1en and IL13Rα2 c1.1en +ΔHER2CD5 CAR-T cell number following 48h treatment with the indicated concentration of T-DM1 (n=6). Positive CAR-T expressing cells were counted using Precision Count Beads. A representative flow cytometry plot is shown on the right. Percentage (**D**) and MFI (**E**) of IL13Rα2 c1.1en and IL13Rα2 c1.1en +ΔHER2CD5 CAR-T cell following treatment with 2.5µg/ml of T-DM1. Cohorts are shown as mean ± SEM. Statistical analysis was done by paired t-tests. *P < 0.05, **P < 0.01, ***P < 0.001, ****P < 0.0001 and not significant (ns) P > 0.05.

### Generation and validation of IL13Rα2 c1.1^en^ multi-armoured CAR

Our findings demonstrate that each individual armoured module functions effectively within the context of the IL13Rα2 c1.1^en^ CAR construct. However, combining all modules into a single transgene cassette poses a significant challenge due to size constraints. Viral delivery systems are typically limited to payloads of approximately 8-9kb, and larger constructs result in diminished expression efficiency that correlates with their increased size^33^. To address this limitation and maximise transduction efficiency, we developed an optimised two-step viral production protocol. This approach involves creating an intermediate viral producer cell line pseudotyped with the vesicular stomatitis virus envelope (VSV-G). This intermediate line is then used to establish a stable viral producer cell line that generates viruses incorporating the RD114 envelope protein, which has been shown to have superior transduction ability in primary human lymphocytes^34,35^ (Supplementary Figure S4A). We then tested this protocol using IL13Rα2 c1.1^en^ Arm CAR, a bicistronic γ-retroviral vector encoding IL13Rα2 c1.1^en^, DHER2^CD5^, dsFvTBRII/CCR and IL12^inv^ all separated by a self-cleaving 2A peptide and transcribed from a single promoter (Supplementary Figure S4B). Upon transduction of autologous T cells with this construct, we observed remarkably efficient expression of each individual component. Notably, and unexpectedly, all components were expressed at a near-perfect 1:1 ratio throughout the entire construct (Supplementary Figure S4B). Testing the IL13Rα2 c1.1^en^ Arm CAR and IL13Rα2 c1.1^en^ CAR using GBM cell lines revealed that the armoured CAR demonstrated superior cytotoxic activity compared to the CAR alone and single armoured CARs (Figure 5A and Supplementary Figure S4D). Notably, we detected significantly higher levels of IFNγ production in the IL13Rα2 c1.1^en^ Arm cells, suggesting that the IL12^inv^ component remained functional (Supplementary Figure S4E). We confirmed this by measuring NK cell pSTAT4 levels, which showed that IL-12 was indeed present and functional (Supplementary Figure S4F).

**Figure 5:**
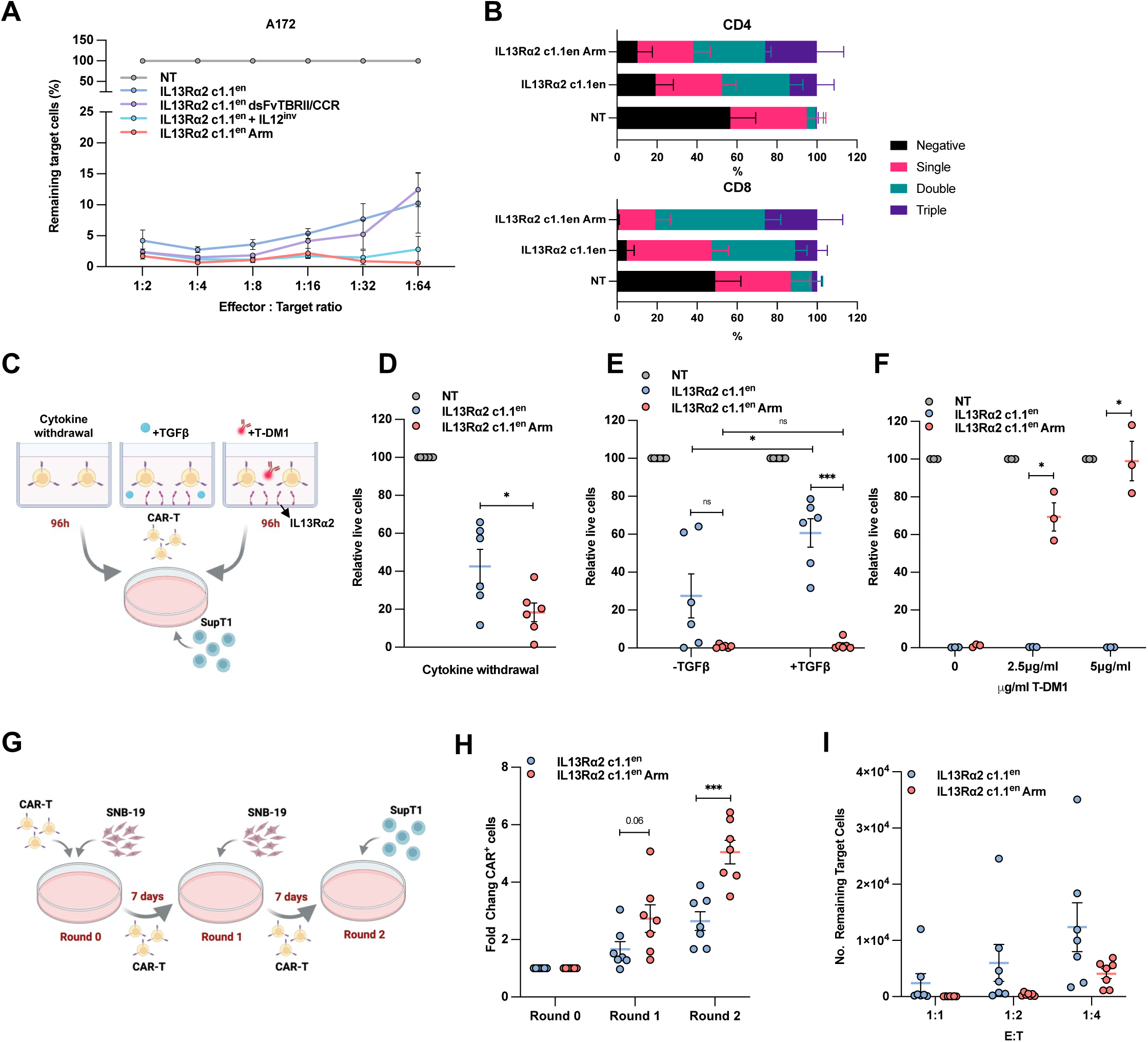
IL13Rα2 c1.1en Arm CAR shows resistance to TGFβ1 and enhanced cell persistence in vitro. Viable target A172 cells following co-culture with IL13Rα2 c1.1en or IL13Rα2 c1.1en Arm CAR-T cells and single armoured CAR-T cells at the indicated E:T ratio quantified by flow cytometry. The percentage of live target cells was calculated by normalizing to the NT group. **B.** Co-expression of TIM3, LAG3 and PD-1 on IL13Rα2 c1.1en and IL13Rα2 c1.1en Arm CAR T cells within CD4⁺ and CD8⁺ subsets after 3-day co-culture with IL13Rα2-expressing targets. **C.** Schematic representation of the experimental layout. In brief, CAR-T cells were cultured in a cytokine-deprived medium or in the presence of 1ug/ml IL13Rα2 immobilised protein with or without TGFβ1 or T-DM1 for 96h. A fixed volume of cells was then transferred to a plate containing SupT1-IL13Rα2 to measure their cytotoxic activity. Following another 96h, viable target cells were quantified by flow cytometry (**D**,**E**,**F**). **G.** Schematic representation of the experimental layout. In brief, CAR-T cells were co-cultured with SNB19 for 7 days (round 0) before being counted and re-seeded on fresh SNB19 for an extra 7 days (round 1). The cells were then counted, normalised and cultivated with SupT1-IL13Rα2 target cells for 48 hours (round 2). The fold increase of CAR-positive cells after each round of replating is shown in (**H**). The cytotoxic activity at the indicated E:T ratio following re-plating after round 1 is shown in (**I**). Cohorts are shown as mean ± SEM. Statistical analysis was done by paired t-tests. *P < 0.05, **P < 0.01, ***P < 0.001, ****P < 0.0001 and not significant (ns) P > 0.05. **C**,**G** were created using BioRender.com

T cell exhaustion poses a major obstacle to long-term CAR-T efficacy, marked by dysfunctional states with reduced proliferation, impaired anti-tumor functions, and co-upregulation of inhibitory receptors like TIM3, LAG3, and PD-1^36^. We therefore assessed these markers via co-expression profiles in transduced cells. After 3-day co-culture with IL13Rα2-expressing targets, TIM3/LAG3/PD-1 expression was evaluated on IL13Rα2 c1.1^en^ and IL13Rα2 c1.1^en^ Arm CAR T cells. In CD4⁺ and CD8⁺ subsets, stimulated CAR T cells showed elevated exhaustion markers versus NT cells; most were double-positive, with triple-positive (TIM3⁺LAG3⁺PD-1⁺) cells limited. Distributions were comparable between IL13Rα2 c1.1^en^ and Arm CAR T cells, indicating our armored elements do not promote terminal exhaustion (Figure 5B).

To evaluate the enhanced capabilities of the IL13Rα2 c1.1^en^ Arm compared to the standard IL13Rα2 c1.1^en^ CAR, we conducted a comprehensive series of stress tests (Figure 5C). Our initial assessment focused on the advantages conferred by the CCR module under cytokine-deprived conditions. Transduced autologous T cells were cultured without cytokines for 96 hours, after which we measured their killing activity against SupT1 IL13Rα2 target cells. The results clearly demonstrated that IL13Rα2 c1.1^en^ Arm-modified T cells exhibited superior expansion, survival, and target cell elimination efficacy in this nutrient-deprived environment compared to their standard CAR counterparts (Figure 5D). We then examined the cells’ killing capacity following antigen stimulation in the presence of TGF-β by culturing CAR-T cells on plates coated with IL13Ra2 protein and TGF-β for 96 hours before assessing their cytotoxic activity. The IL13Rα2 c1.1^en^ Arm consistently outperformed the IL13Rα2 c1.1^en^ CAR under these conditions (Figure 5E). To test the regulatory capability of the HER2 module, we cultured the cells in the presence or absence of T-DM1. Following T-DM1 exposure, the cells lost their ability to kill target cells and secrete IFNγ, demonstrating effective regulation through the safety switch (Figure 5F and Supplementary Figure S4G). To further assess the superior performance of the IL13Rα2 c1.1^en^ Arm CAR, we conducted a stress test using SNB19 glioblastoma cells. CAR-T cells were co-cultured with SNB19 cells for 7 days, then transferred to freshly plated SNB19 cells for an additional 7 days. Following this extended challenge, CAR-T cells were normalized and co-cultured with SUPT1-IL13Rα2 cells to measure their killing activity (Figure 5G). The results demonstrated that the IL13Rα2 c1.1^en^ Arm significantly outperformed the IL13Rα2 c1.1^en^ CAR, even after prolonged exposure to tumour cells (Figure 5H,I). These comprehensive tests highlight the advantages of our armoured CAR design, demonstrating improved persistence, killing efficiency, and maintained functionality under various challenging conditions that mimic the hostile tumour microenvironment. To further evaluate the IL13Rα2 c1.1^en^ Arm CAR-T cells in a more physiologically relevant manner, we used a well characterised GBM tumour model by orthotopically injecting U87 cells, modified to stably overexpress IL13Rα2, in immunodeficient mice^37^. Following 8 days of engraftment, mice were grouped based on similar bioluminescence signals and intracranially injected with 6.25*10^4^ CAR-T cells (Figure 6A). Remarkably, while IL13Rα2 c1.1^en^ CAR-T cells failed to eradicate tumors in 3 out of 5 mice, complete tumour eradication was achieved in all 5 mice treated with IL13Rα2 c1.1^en^ Arm CAR-T cells (Figure 6B,C). Importantly, the body weight of the animals remained stable throughout the study, underlining the safety profile of our CAR-T construct (Figure 6D). We also assessed IL13Rα2 c1.1^en^ Arm CAR-T cells safety and efficacy in a subcutaneous U87-IL13Rα2 xenograft model. Tumors reaching 200 mm^3^ were treated with intravenous CAR-T cells; after 30 days, IL13Rα2 c1.1^en^ Arm CAR-T outperform non-armored controls, with persistent circulating IL13Rα2 c1.1^en^ Arm CAR-T cells detectable by FACS (non-armored undetectable) (Figure 6E,J). Sustained tumor control was also observed at lower CAR-T doses (Supplementary Figure 5A). To evaluate safety, peripheral IFN-γ was measured in blood from treated animals. Animals receiving IL13Rα2 c1.1^en^ Arm CAR-T cells showed higher IFN-γ levels than those treated with IL13Rα2 c1.1^en^ CAR-T alone, where IFN-γ was undetectable, aligning with limited CAR-T detection and higher tumor burdens in the latter (Figure 6K and Supplementary Figure 5B). Notably, despite detectable IFN-γ in the IL13Rα2 c1.1^en^ Arm cohort, no overt systemic toxicity occurred, as body weights remained comparable to CAR-alone and NT controls without decline. These findings collectively highlight the superior anti-tumour efficacy of our IL13Rα2 c1.1^en^ Arm CAR-T cells in a clinically relevant GBM model, while also demonstrating the potential for precise control of CAR-T cell activity to enhance safety.

**Figure 6:**
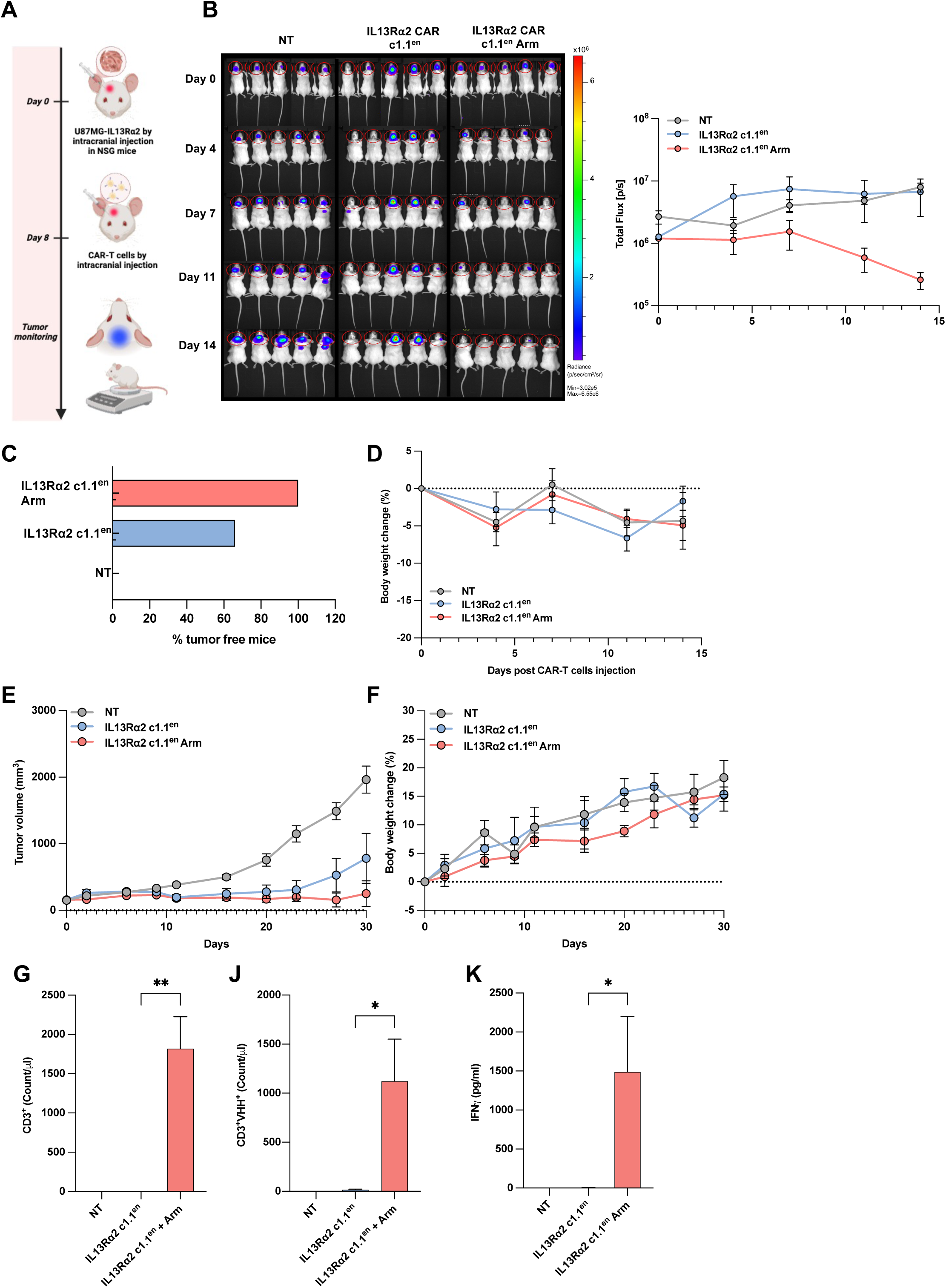
IL13Rα2 c1.1en Arm CAR shows enhanced cell persistence in vivo. **A.** Design of the orthotopic in vivo experiment. (created using BioRender.com) **B.** Bioluminescence (BLI) images and values at day 0, 4, 7, 11 and 14 after untransduced (NT), IL13Rα2 c1.1en or IL13Rα2 c1.1en Arm CAR-T cells injection. **C.** Bars represent the percentage of tumor-free mice at day 14. **D.** Body weight at day 0, 4, 7, 11 and 14 after untransduced (NT), IL13Rα2 c1.1en or IL13Rα2 c1.1en Arm CAR-T cells injection. Tumor volume (**G**) and body weight (**H**) measurements of NSG mice subcutaneously injected with U87-IL13Rα2 target cells, which were allowed to engraft for 8 days before a 5×106 dose of non-transduced T or CAR-T cells was intravenously injected via the tail vein. **I,J.** Number of hCD3+ and HA+ cells quantified by FACS analysis at sacrifice (day 30). **K.** Quantification of IFN-γ levels in plasma samples collected from mice at day 30 post-treatment. IFN-γ concentrations were measured by ELISA. Cohorts are shown as mean ± SEM. Statistical analysis was done by paired t-tests. *P < 0.05, **P < 0.01, ***P < 0.001, ****P < 0.0001 and not significant (ns) P > 0.05.

### Validation of the IL13Rα2 c1.1^en^ multi-armoured CAR for clinical purpose

Given the complexity of our CAR-T construct, we aimed to evaluate the feasibility of manufacturing IL13Rα2 c1.1^en^ Arm CAR-T cells within a Good Manufacturing Practice (GMP) workflow. We activated sorted CD3^+^ T cells from two healthy donors using T Cell TransAct™. Forty-eight hours later, these cells were transduced with clinical-grade IL13Rα2 c1.1^en^ Arm viral supernatant (titer 4.89×10^8^ TU/ml) at varying multiplicities of infection (MOI). The cells were then cultured using the G-Rex cell culture platform, which utilises gas-permeable membrane technology to enhance oxygen delivery and nutrient exchange^38^, in the presence of IL7/IL15 and an optimized serum-free cell culture medium (Figure 7A). Notably, we observed a dose-dependent transduction efficiency, achieving a good level of CAR expression (>20%) even at the lowest MOI tested, with peak expression occurring five days post-transduction (Figure 7B). According to current FDA recommendations, the maximum allowable vector copy number (VCN) is five copies per transduced cell for virally transduced cell therapy products^39^. Our analysis of VCN seven days post-transduction confirmed that this limit was not exceeded by any of the MOIs tested, demonstrating the construct’s compatibility with GMP standards and regulatory guidelines (Figure 7C). The expansion and propagation of the cells were monitored for up to 10 days following T cell isolation. Results showed a significant increase in cell numbers, with up to 30-fold expansion and an average total T cell count reaching 10^9^ cells (Figure 7D). Further analysis of T cell subsets revealed a shift in the composition over time. There was a progressive decrease in T memory stem (Tscm) cells, accompanied by an increase in effector memory T cells (Tem cells). Interestingly, the percentage of central memory (Tcm) cells, a population with self-renewal capability, better persistence and antitumor immunity^40^, remained stable throughout the expansion period (Figure 7E). On the final day of manufacturing (day 10), the CD4/CD8 ratio was approximately 1:4 (Figure 7F). The cells exhibited expression of Lag3 and Tim3, with low expression of PD1 (Figure 7G). To validate the functionality of our CAR-T cells at the end of the manufacturing process, we assessed their ability to kill target cells. The data demonstrated that the cells maintained significant killing activity even after the 10-day manufacturing period (Figure 7H). Together, these data demonstrate that IL13Rα2 c1.1^en^ Arm CAR-T cells can be manufactured using a GMP adaptable process while retaining their cytotoxic capabilities, suggesting their potential effectiveness in targeting cancer cells upon administration.

**Figure 7:**
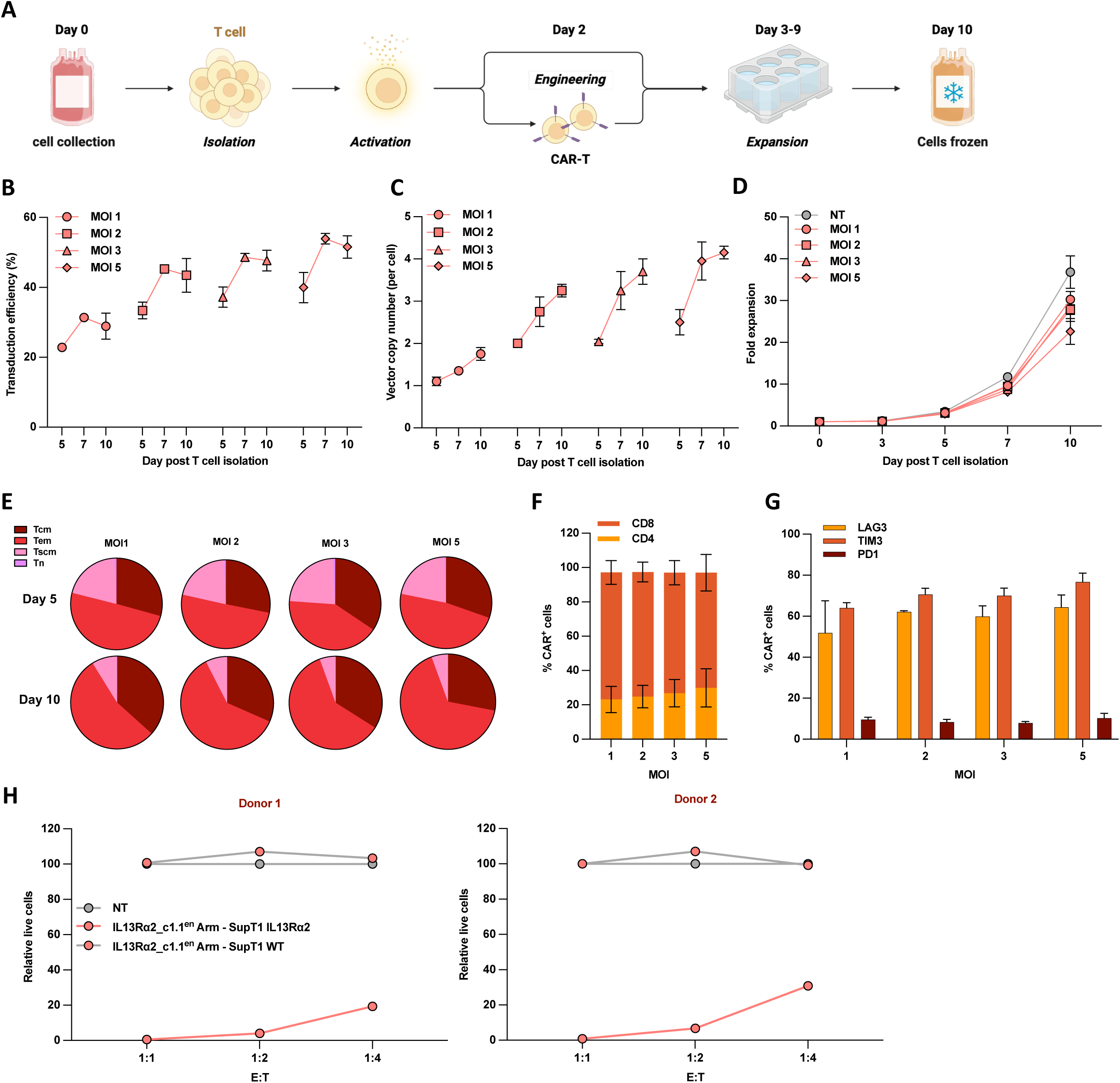
Validation of the IL13Rα2 c1.1enmulti-armoured CAR (NS007) for clinical use. **A.** Schematic representation of the manufacturing protocol created with biorender.com. Transduction efficiency (**B**) and vector copy number (VCN) (**C**) at day 5, 7 and 10 post T cell isolation at the indicated multiplicity of infection (MOI). **D.** Fold change in T cell expansion over a period of 10 days**. E.** Naïve (Tn), central memory (Tcm), effector memory (Tem) and memory stem cells (Tscm) T-cell subsets were identified following gating on CD45RA and CCR7 expression. Ratio of CD4:CD8 (**F**) and expression of exhaustion markers (**G**) in CAR-T positive cells 10 days post T cell isolation. **H**. Viable target SupT1-IL13Rα2 and SupT1 cells following co-culture with GMP grade manufactured IL13Rα2 c1.1en Arm CAR-T cells at the indicated E:T ratio quantified by flow cytometry. The percentage of live target cells was calculated by normalizing to the NT group. Cohorts are shown as mean ± SD of two independent healthy donors.

## DISCUSSION

Recent clinical studies have demonstrated promising results for CAR-T cell therapy in GBM patients. However, the efficacy of this approach has been constrained by limited CAR-T cell persistence and the immunosuppressive nature of the TME. To address these challenges, we describe here a novel, multi-functional CAR-T construct within a single cassette.

IL13Rα2 represents an exciting molecular target for cell therapy, primarily due to its specific expression in GBM and its presence in cancer stem cells. Most current efforts, including clinical trials, have utilised the IL13 binder known as Zetakine to target IL13Rα2. However, a significant challenge with using IL-13 Zetakine is its lack of selectivity, as it also binds to IL13Rα1. To address this cross-reactivity, researchers have developed a mutated form (E13Y) for CAR-T cell preparation, although this approach has shown limited efficacy. Subsequent enhancements included the incorporation of different co-stimulatory molecules that increased efficacy^41,42,43,44^. These findings have undoubtedly contributed to the advancement of IL13Rα2 cell therapies for GBM. Our studies corroborated these conclusions while also revealing a novel high-throughput approach to identify new binders for IL13Rα2-targeted therapy. Through a humanization process, we successfully identified a new enhanced IL13Rα2 binder that demonstrates high specificity for IL13Rα2. The approach enabled the generation of a highly tractable single-domain antibody with enhanced selectivity for IL13Rα2, reduced immunogenicity, and excellent biophysical properties, making it more suitable for development as a CAR targeting domain for the treatment of GBM.

Two recent clinical studies involving patients with recurrent glioblastoma treated with locally infused CAR-T cells demonstrated tumour reductions in nearly all participants, with substantial regressions observed within 24-48 hours post-infusion. However, the majority of these responses proved to be transient, with tumour recurrence noted in subsequent follow-ups^45,10^. One proposed solution to this challenge is the administration of repeated doses of CAR-T cells. While this approach has been shown to be safe in a cohort of 65 patients, it presents significant limitations in terms of feasibility, primarily due to the complexities and costs associated with cell manufacturing^11^. To address these limitations, we opted for an alternative strategy focused on enhancing the CAR-T cells’ ability to secrete or express factors that can counteract the immunosuppressive nature of the tumour microenvironment. Indeed, we focused our attention on two key factors: TGF-β and IL-12.

TGF-β is known to induce Treg-like cell conversion and PD-1-dependent T cell exhaustion, significantly inhibiting the proliferation and activity of CAR-T cells both *in vitro* and *in vivo*^46,47,14^. To counteract this effect, we developed a novel disulfide-stabilized Fv antibody targeting TGF-β receptor II, capable of blocking the downstream activation of TGF-β signalling. We further innovated by coupling this antibody with the α and β chains of the GM-CSF receptor to generate a constitutive active intracellular signal, leveraging the disulfide bond for this purpose. The expression of this combined module led to remarkable results, with cells persisting *in vitro* for over a month. These engineered cells demonstrated the ability to undergo multiple rounds of expansion while maintaining impressive *in vitro* cytotoxicity even after TGF-β treatment. Notably, our experiments revealed that the β/α orientation of the GM-CSF receptor chains produced a more intense signal compared to the α/β configuration, as evidenced by enhanced STAT5 activation. This finding suggests that our module not only offers the advantage of being cytokine or stimulus-independent compared to other approaches^48^, but also allows for modulation based on CAR architecture or when a more dampened STAT5 intracellular activation is desired. While the enhanced proliferation capacity of these cells could potentially lead to concerns about lymphoproliferative syndromes, our in vitro data suggest that the presence of the GM-CSF CCR alone is not sufficient to cause such complications, as the cells eventually undergo natural cell death. Notably, a similar approach using a signalling cytokine receptor that triggers the IL-7 signalling axis (C7R) has been recently shown to be well-tolerated in young patients with CNS tumours^49,50^. This suggests a low risk for adverse events if our CAR T-cell and CCR combination strategy were to be employed for GBM treatment.

IL-12 is a crucial cytokine that plays a significant role in the regulation of immune responses by mainly enhancing the cytotoxic activity of NK and T cells through increased production of IFNγ^51,52^. Given its powerful antitumor function, there has been considerable interest in using IL-12 as an adjuvant in cell therapy and immunotherapy^53^. However, due to its potency being coupled with high toxicity, recent efforts to develop more tolerable versions of the cytokine have been explored. Attempts to regulate the expression of a secretable IL-12 under an inducible NFAT promoter in adoptively transferred T-cell therapy still resulted in unwanted toxicities due to the systemic distribution of the cytokine or a partial leakage of the NFAT gene regulation system^54^. To address this issue, many research groups have explored membrane-bound or -tethered approaches, sometimes coupled with locoregional delivery of CAR-T cells, to limit IL-12 distribution^55,56,57^. While these strategies have demonstrated improved safety profiles, they may potentially limit the full anticancer potential of IL-12 by restricting its ability to activate other cell types. To avoid these limitations, we pursued a different approach to harness the benefits of IL-12 while mitigating its systemic toxicity. Indeed, we leveraged the heterodimeric nature of the IL-12 protein, which is composed of p35 and p40 subunits that together form the biologically active IL-12p70. We created a single-chain form that expresses the p35 subunit in front of the p40 (IL12^inv^), a configuration known to reduce the protein’s biological activity. Through experimentation with various linker lengths separating the two subunits, we discovered that a 7-amino acid linker most effectively decreased the biological activity of the construct. Our data suggest that this reduction is primarily due to a lower affinity for the IL-12Rb1 receptor. While a more comprehensive analysis is needed, our current interpretation is that inverting the two subunits either inhibits the correct folding of the protein or causes the p35 subunit to mask the p40 region responsible for receptor binding. Nevertheless, given that p40 binding to the receptor is the initial and limiting step in IL-12 signalling^27^, our inverted scIL12 exhibits significantly reduced activation potency. This novel approach results in a protein that still retains all the IL-12’s beneficial properties but with a much lower risk of systemic toxicity. In conclusion, by fine-tuning the structure and activity of IL-12, we created a safer version of this potent cytokine for use in cancer immunotherapy, potentially overcoming the limitations of previous strategies while maintaining its broad immunostimulatory effects.

CAR-T therapy is generally considered safe, but severe side effects such as cytokine release syndrome (CRS) and immune effector cell-associated neurotoxicity syndrome (ICANS), though rare, can still occur. To address these risks, safety switches have become a crucial component in cell therapy development, offering a mechanism to control or eliminate engineered cells in case of severe toxicities. While various approaches are being explored, the most common strategies aim to eliminate CAR-T cells entirely. Among these approaches, the inducible caspase-9 suicide gene system (iCas9), ganciclovir-activated switches, and antibody-dependent cell-mediated cytotoxicity (ADCC) switches are the most frequently utilised. The iCas9 method has shown promising results in clinical trials^58,59^, but its clinical application is limited by the lack of regulatory approval for its inducer drugs (AP1903 or AP20187) in cell therapy. Ganciclovir-activated switches, derived from non-human sequences, pose potential immunogenicity risks, potentially inducing both humoral and cellular immune responses against CAR-T cells. Given these limitations, we focused our efforts on developing an ADCC switch. We engineered a truncated version of HER2, retaining only domain IV, which binds to T-DM1, a safe and commercially available drug. Additionally, to enhance the internalisation efficiency of the drug, we replaced the HER2 transmembrane domain with that of CD5. Our preliminary results demonstrate almost complete clearance of CAR-T cells within 48 hours *in vitro*. Furthermore, this T-DM1-based switch is compatible with established GBM delivery routes like Ommaya reservoirs or intra-arterial infusion, which bypass the blood-brain barrier for targeted intracranial drug access without systemic toxicity. The efficacy of our approach, achieving approximately 85% elimination, is comparable to clinically used iCasp9-based strategies^60^. This offers a potentially safer and more clinically applicable method for controlling CAR-T cell activity, addressing key challenges in current safety switch technologies.

In conclusion, our research represents a promising advancement in CAR T-cell therapy. Despite recent encouraging outcomes, CAR-T cell therapy for solid tumours remains a significant challenge, primarily due to the transient nature of treatment responses observed in many clinical trials. These challenges are largely attributed to factors within the hostile TME. Our ability to integrate multiple functional enhancements within a single construct that can still be manufactured marks a substantial step forward in the field. Moreover, our findings extend beyond treating GBM, as they can serve as standalone armoured modules for various CAR-T constructs targeting different solid tumours. Demonstrating the feasibility of manufacturing such a complex construct opens the door to incorporating diverse combinational elements within a single CAR-T construct, allowing for tailored treatment strategies based on specific cancer TMEs and antigens. Indeed, our data offers hope for more effective and safer treatments for patients with GBM and other difficult-to-treat cancers.

### Limitations of the study

Our research underscores the value of leveraging computational and in silico data in the design of enhanced VHH CAR constructs. Although we demonstrate that this approach could potentially reduce immunogenicity and enhance binding specificity, it is important to note that our conclusions are derived from *in vitro* studies. Furthermore, while our multi-modular strategy shows promise, it may also present limitations that require further investigation. We have successfully demonstrated the safety and tolerability of our multi-armoured CAR-T design in both *in vitro* cell culture systems and mouse models. However, it is crucial to recognize that syngeneic mice and especially immunocompromised mouse models might differ substantially from the actual human response and tolerability to cytokines such as IL-12. This discrepancy highlights the need for caution when extrapolating our findings to human applications. Finally, we demonstrated the feasibility of manufacturing our product, but it is important to note that this process was conducted using specific conditions and may require further optimization for large-scale production or clinical applications.

## RESOURCE AVAILABILITY

### Lead contact

Requests for further information and reagents should be directed to and will be fulfilled by the Lead Contacts, Maurizio Mangolini and Shimobi Onuoha

### Materials availability

All unique reagents generated in this study are listed in the manuscript and available from the lead contacts upon reasonable requests and a completed Materials Transfer Agreement.

### Data and code availability

Transcriptomic data and accession codes will be made available upon manuscript’s acceptance.

## Acknowledgements

We acknowledge the contribution of many people who contributed to this work either through useful discussions or the provision of reagents, including Agnieszka Andres and Dr Jun Li.

## Author contributions

Conception and experimental design: MM, SO, SS, SC, BM

Computational data generation: PS, SO, AR, MG

Experimental data generation: MM, ES, DP, SS, RK, YY, LS, HW

Analysis and interpretation of data: MM, SO, SS, SC, BM

Writing: MM, SO, SS, SC, PS

## METHOD DETAILS

### Primary cells and cell culture

Peripheral Blood Mononuclear Cells (PBMCs) were isolated from Leukocyte Reduction System (LRS) cones from consented platelet apheresis donors (NHSBT, Cambridge, UK) by density centrifugation via Ficoll-Paque PLUS (GE-Healthcare; GE17-1440-03) according to manufacturer protocol. Isolated PBMCs were cultured in complete RPMI (GIBCO, ThermoFisher Scientific, Winsford, UK) supplemented with 10% Fetal Calf Serum (FBS) (Labtech, Heathfield, UK), 2mM GlutaMAX (Gibco; 35050061), IL-7 and IL-15 at ng/mL (Miltenyi Biotec, Germany).

Natural Killer (NK) cells were generated from isolated PBMCs using the CellXVivo Human NK Cell Expansion Kit (Bio-techne, R&D System, Minneapolis, US) according to the manufacturer’s instructions.

Mouse splenocytes were isolated from spleens of 6- to 8-week-old C57BL/6 mice (Charles River Laboratories).

HEK-293^GP^ and HEK-293^RD1^^14^ (Biovec Pharma, Quebec City, Canada), Phoenix-ECO (ATCC; ATCC^®^CRL-3214™) SNB19 (DMSZ, ACC 325 Leibniz Institute, Germany) lines were cultured in Dulbecco’s Modified Eagle Medium (DMEM) (GIBCO, ThermoFisher Scientific, Winsford, UK) supplemented with 10% Fetal Calf Serum (FBS) (Labtech, Heathfield, UK) and 2mM GlutaMAX (Gibco; 35050061). HEK-Blue™ TGF-β and HEK-Blue™ IL-12 Cells (InvivoGen, Toulouse, France) lines were cultured in complete DMEM supplemented with a selection of antibiotics according to the manufacturer’s instructions. SupT1 (ATCC; ATCC^®^CRL-1942™) and Jurkat (Clone E6, ATCC; ATCC^®^TIB-152™) cells were cultured in complete RPMI 1640 (GIBCO, ThermoFisher Scientific, Winsford, UK) supplemented with 10% FBS and 2mM GlutaMAX (Gibco; 35050061).

### Retroviral production and Transduction

For amphotropic viral production, HEK-293GP cells were transiently transfected with a VSV-G envelope expression plasmid and a retroviral cDNA construct (ratio 1:4). The transfection was carried out with GeneJuice (Millipore; 70967-4) according to the manufacturer’s guidelines (ratio DNA:GeneJuice 1:2.5). After 72 h, the cell supernatant was filtered through a 0.45μM filter and the viral supernatant was added to HEK-293^RD114^ cells. Transduction was performed by spinoculation (1500 × *g*, 1 h at 32 °C) with the addition of 10μg/ml Polybrene (INSIGHT Biotechnology). Viral supernatant from HEK-293^RD114^ cells was then collected at different time points when cell confluency reached ∼100%, snap frozen and stored at -80 °C for subsequent titration and use. Isolated PBMCs were activated using 10ng/ml IL-7 and IL-15 and 1:100 T Cell TransAct™ (Miltenyi Biotec). 48 hours post activation, 1×10^6^ PBMCs were plated on Retronectin (Takara Clonetech; T100B) on 6 well plates (Corning; 351146) with viral supernatant and spun for 1500×*g*, 1 h at 32 °C.

For ecotropic viral production, Phoenix-ECO cells were transiently transfected with the pCL-ECO packaging vector and a retroviral cDNA construct. After 72 h, the cell supernatant was filtered through a 0.45μM filter and frozen. Isolated murine splenocytes were activated using 50ng/ml of aCD3 (Miltenyi Biotec; 130-093-387)/ aCD28 (Miltenyi Biotec; 130-093-375) and 50U/ml of mIL-2 (Miltenyi Biotec; 130-120-330). 24 hours post activation, cells were plated on Retronectin (Takara Clonetech; T100B) on 6 well plates (Corning; 351146) with viral supernatant and spun for 1500×*g*, 1h at 32 °C

### Flow Cytometry

Flow cytometry was performed using a MACSQuant 10 flow cytometer (Miltenyi Biotec, Germany). All labelling was conducted at room temperature (RT) for 15 minutes in subdued light with antibodies diluted in PBS+2% FBS. The antibody used are: aCD3-PECy7 (BioLegend; 300420), aCD2-PE (BioLegend; 300208), aHA-APC (BioLegend; 901524), aCD56-FITC (BioLegend; 318304), aCD8-FITC (BioLegend; 344704), aProtein L (Cell Signalling, 29480L), pSTAT4-AF647 (BD Biosciences, Oxford, UK 558137) apSMAD2/3-APC (BD Biosciences, Oxford, UK 562586), aHER2-FITC (R&D System, Minneapolis, US FAB9589G).

IL13Rα2 CAR was detected using MonoRab™ Rabbit Anti-Camelid VHH Cocktail iFluor 647 (GenScript, Oxford,UK A02019-200).

Intracellular staining was performed using BD Phosflow™ Fix Buffer I and BD Phosflow™ Perm Buffer III (BD Biosciences, Oxford, UK) according to the manufacturer’s protocol.

### Cytotoxicity assay

Transduced cells were co-cultured with target cells at various effector:target (E:T) ratios, with target cells remaining constant at 5×10^4^. Target cells are counted via flow cytometry using CountBright Absolute Counting beads (Life Technologies; C36950) and cytotoxicity was calculated as a percentage by normalising to the number of target cells recovered from co-cultures with non-transduced (NT) T cells.

### Plate bound and re-stimulation assay

Normalised T cells were cultured in 24-well plates previously coated overnight with 1ug/ml IL13Rα2 protein and 1ug/ml TGF-β. 96h later cells were then co-cultured with SUPT1 target cells and cytotoxicity was measured after 96h.

CAR T cells were seeded onto SNB19 cells and maintained for 7 days. Cells were harvested and counted (Countess 3 Cell Counters, ThermoFisher Scientific, Winsford, UK). Cells were then transferred to freshly plated SNB19 cells for repeat stimulation. 7 days later, T cells were harvested, counted and finally co-cultured with SUPT1 target cells for cytotoxicity analysis at the indicated E:T ratio.

### Cell-Free Protein Expression

Proteins were produced using a cell-free expression system, specifically the PUREfrex® 2.0 system (GeneFrontier Corporation), according to the manufacturer’s protocol. Synthesized constructs were derived from gBlocks® gene fragments (Integrated DNA Technologies), designed to include a T7 promoter for transcription and a polyadenylation (poly A) tail for stability. Each construct encoded a His-tag for subsequent purification. After transcription and translation, the reaction mixtures were subjected to Ni-NTA affinity chromatography for protein purification. The eluted proteins were concentrated and buffer exchanged using Amicon Ultra centrifugal filters (MilliporeSigma) before downstream analyses. Protein purity and yield were assessed via SDS-PAGE and Bradford assay.

### Mammalian Expression

A separate set of proteins was produced in a mammalian expression system using the pcDNA3.1(+) vector (Thermo Fisher Scientific). Constructs were designed to encode target proteins fused with an Fc tag for improved stability and purification. Transient transfections were performed in CHO-S cells (Thermo Fisher Scientific) cultured in CD CHO medium supplemented with 8 mM L-glutamine and 1% HT supplement (Gibco). Transfections were carried out using polyethylenimine (PEI) at a DNA:PEI ratio of 1:3. Following transfection, cells were incubated for 5–7 days in a shaking incubator at 37°C with 5% CO₂ and 80% humidity.

The proteins were harvested from the culture supernatant and purified using Protein A affinity chromatography (GE Healthcare) under gravity flow. Eluted proteins were neutralized immediately with 1 M Tris-HCl (pH 8.0) and subjected to buffer exchange into PBS using Amicon Ultra centrifugal filters. The purity of the purified proteins was assessed using SDS-PAGE, and concentrations were determined using a NanoDrop spectrophotometer (Thermo Fisher Scientific). Proteins were stored at −80°C until further use.

### Binder Discovery and NGS sequence analysis

NGS data were obtained from the MiSeq sequencing platform as two fastq files, one for forward and one for reverse reads. These files were processed with an in-house python script that merges forward and reverse reads assuming an overlapping region (which is always present for short proteins like VHHs) to obtain the full sequence of the VHHs. The script further counts the number of times each unique sequence is observed in the raw files (i.e., the number of reads per unique sequence). Sequences were in silico translated to amino acids, and their count score and corresponding frequency were used to sort the observed sequences from most to least frequent, under the assumption that those sequences most abundant in the post-panning library should be those binding most tightly to the target. All VHH sequences observed at least 3 times (to reduce random sequencing errors) were aligned using the AHo numbering scheme as described in ^19^. The aligned CDR1, 2, and 3, were then combined and these joint CDR sequences were used to determine homology groups. Specifically, each of the top 100 sequences by frequency in the library was considered as query, and its homology group contained all sequences with a joint-CDR chemical distance smaller than 94, as calculated from the distance matrix in Supplementary Figure 1a, which corresponds to around 25% CDR sequence identity (this percent varies because e.g., a mutation R to K will have a much smaller chemical distance than one from R to G). The assumption is that all sequences in a homology group should engage the same epitope, given their highly similar CDRs. Overlapping homology groups were merged. Figure 1c reports the percent frequency of the top-ranking sequence used as query on the x-axis, and the sum of the percent frequencies of all sequences in the homology group on the y-axis. Value labels are the ranking of the top sequence in the library, with 0 corresponding to the highest frequency sequence.

### In vivo studies

Animal studies were performed under a UK Home Office–approved project license. Crown Bioscience UK Ltd carried out C57BL/6 animal studies. In brief, 1×10^5^ B16.F10 cells, modified to express GD2, were subcutaneously injected into 6–10-week-old C57BL/6 female mice. Eight days post engraftment 2.5×10^6^ CAR T-cells were injected intravenously and mice were monitored for body weight and tumor volume using a caliper. Mice were randomized one day pre-tumor injection and one day prior to treatment.

5×10^4^ U87 cells, modified to express hIL13Rα2, were inoculated intracranially into NSG mice. On day 7 Tumour engraftment was measured by bioluminescent imaging utilizing the IVIS spectrum system (PerkinElmer, Waltham, MA) after intra-luciferin peritoneal injection and mice were randomized for CAR treatment. The following day, 6.25×10^4^ CAR T-cells were injected intracranially. For the subcutaneous model, mice were subcutaneously injected with 1 × 10⁷ U87-IL13Rα2 cells. CAR-T cells were administered intravenously when tumors reached ∼200 mm³.

### Statistical analysis

Data analyses were performed using GraphPad Prism 10 (GraphPad Software, La Jolla, USA) with paired analyses as indicated. For experiments where more than two groups are compared, statistical analyses were performed using one-way ANOVA followed by two-tail Student t-tests. Statistical annotations were denoted with asterisks as follows: ****P < 0.0001, ***P < 0.001, **P < 0.01, *P < 0.05, and not significant (ns) P > 0.05.

## STAR★METHODS

### KEY RESOURCES TABLE

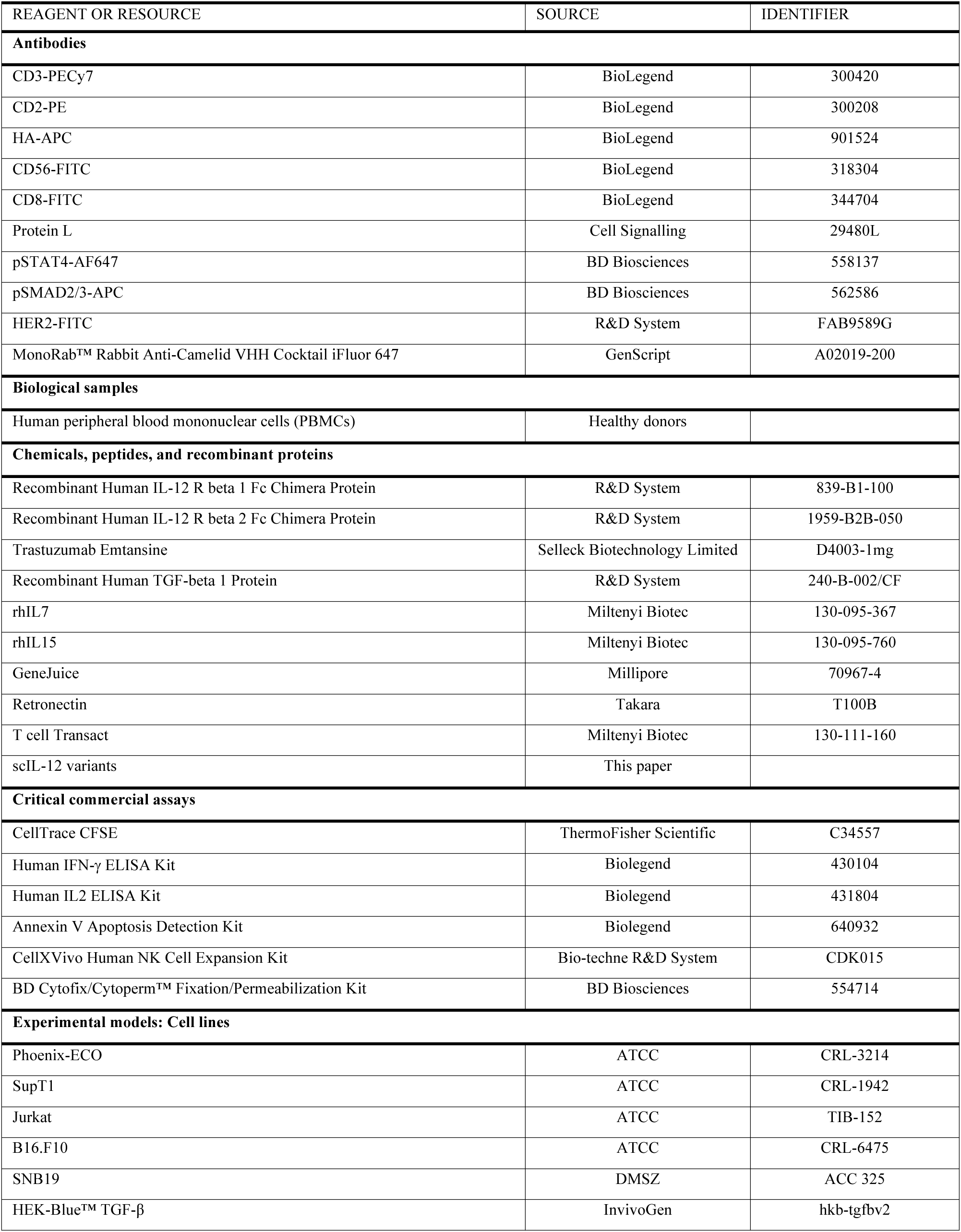

**Supplementary Figure S1:**
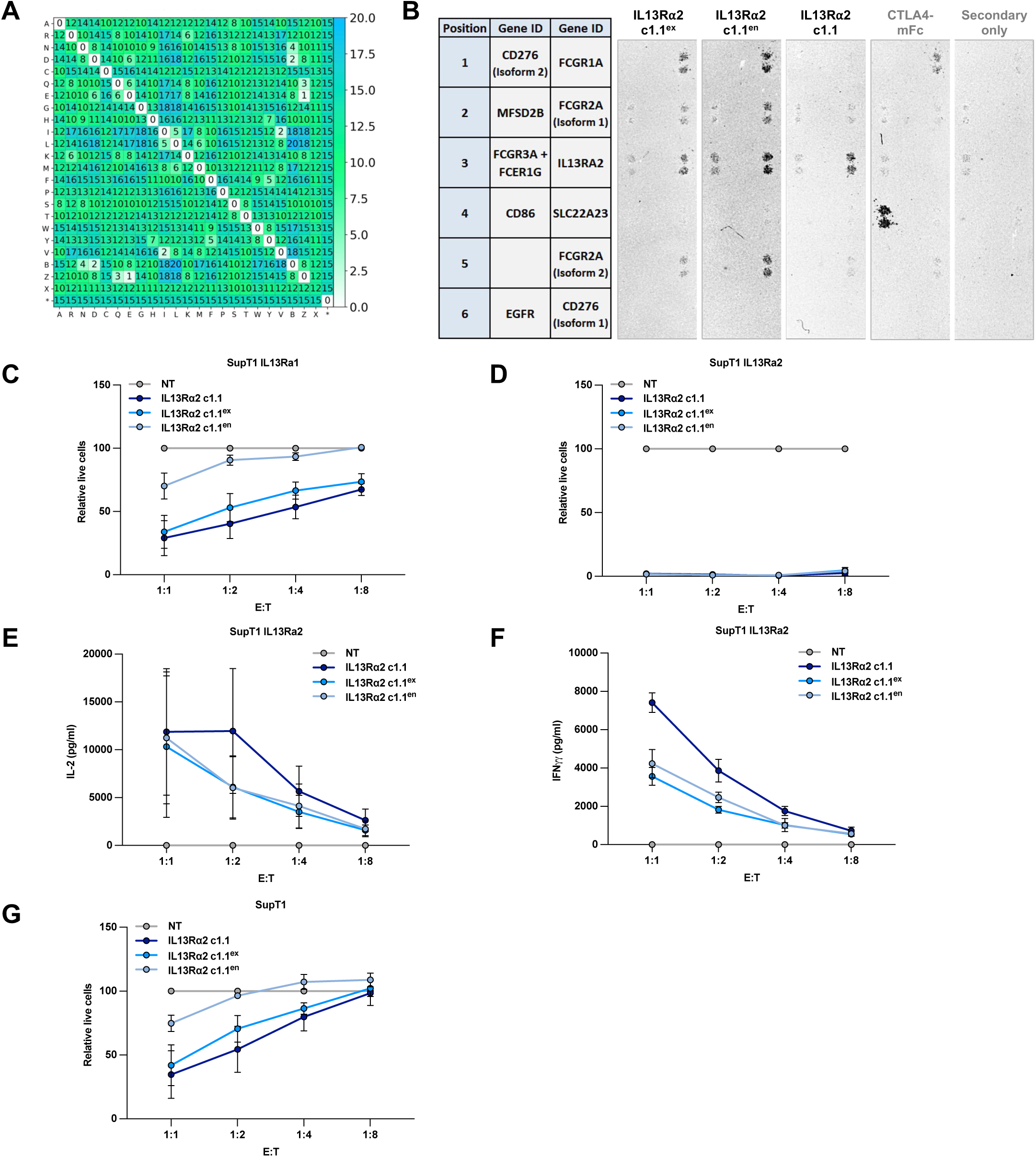
*In vitro* screening identifies lead IL13R⍺2 VHH for CAR-T cells. **A.** Distribution matrix depicting chemical distances between amino acids based on substitution scores**. B.** The IL13R⍺2 c1.1, IL13R⍺2 c1.1ex, and IL13R⍺2 c1.1en VHHs were Fc-conjugated and used to stain an array containing ∼6000 native human membrane proteins. Binding to Fc gamma receptors (e.g., FcGR1A and FcGR2A isoforms) is expected due to the Fc conjugation. The figure shows representative intensity of the staining against selected proteins. Each column corresponds to a different VHH construct, with two dots per protein indicating duplicate spots on the array for technical validation. **C,D.** SupT1 expressing IL13Rα1 (**C**) or IL13Rα2 (**D**) were co-cultured with different ratios of CAR-T cells for 72 hours. Viable target cells were quantified by flow cytometry using CD2 and CD3 expression as exclusion markers. The percentage of live target cells was calculated by normalizing to the NT group. **E,F.** ELISA detection of IL-2 and IFN-γ production by CAR-T cells co-cultured with SupT1-IL13R⍺2 cells at the indicated E:T ratio (relative to Figure 1h). **G**. Quantification of live SupT1 cells following co-culture with CAR-T cells at the indicated E:T ratio for 72h.

**Supplementary Figure S2:**
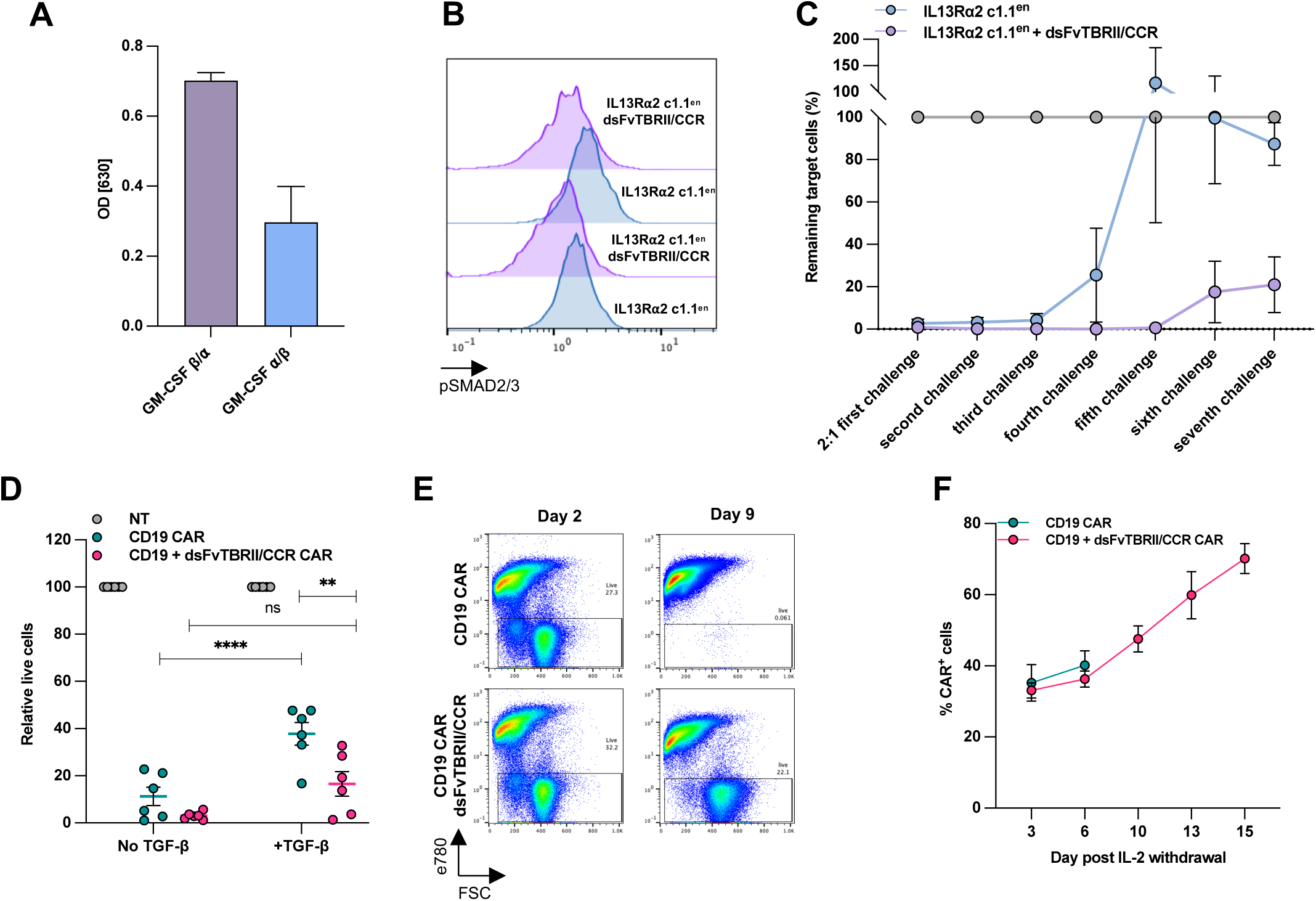
dsFvTBRII/CCR overcomes the inhibitory effects of TGF-b and induces CAR-T cell persistence. **A.** Quantification of the SEAP secreted by HEK-Blue™ TGF-β expressing the dsFvTBRII region fused with different orientations of the GM-CSF receptor chains: alpha-beta and beta-alpha. **B.** Representative FACS histograms of pSMAD2/3 expression following TGFβ stimulation (related to Figure 2E). **C.** T cells expressing IL13R⍺2 c1.1en CAR or IL13R⍺2 c1.1en CAR+dsFvTBRII/CCR were harvested and co-cultured with fresh SNB19 once a week for 7 weeks. Remaining SNB19 cells were quantified using precision counting beads via FACS. **D**. SupT1 expressing CD19 were co-cultured with different ratios of CAR-T cells for 72h with or without 10ng/ml TGFβ. Then, viable target cells were quantified by flow cytometry using CD2 and CD3 expression, and the percentage of live target cells was calculated by normalizing to the NT group. **E.** Representative flow cytometry plot from an individual T cell donor expressing CD19 CAR or CD19 CAR+dsFvTBRII/CCR following cytokine withdrawal at the indicated time point. A total of three independent experiments were conducted, each using lymphocytes from a different healthy donor. **F.** Analysis of CAR-positive cell enrichment over time following cytokine withdrawal measured by flow cytometry (n=3). Cohorts are shown as mean ± SEM. Statistical analysis was done by paired t-tests. *P < 0.05, **P < 0.01, ***P < 0.001, ****P < 0.0001 and not significant (ns) P > 0.05.

**Supplementary Figure S3:**
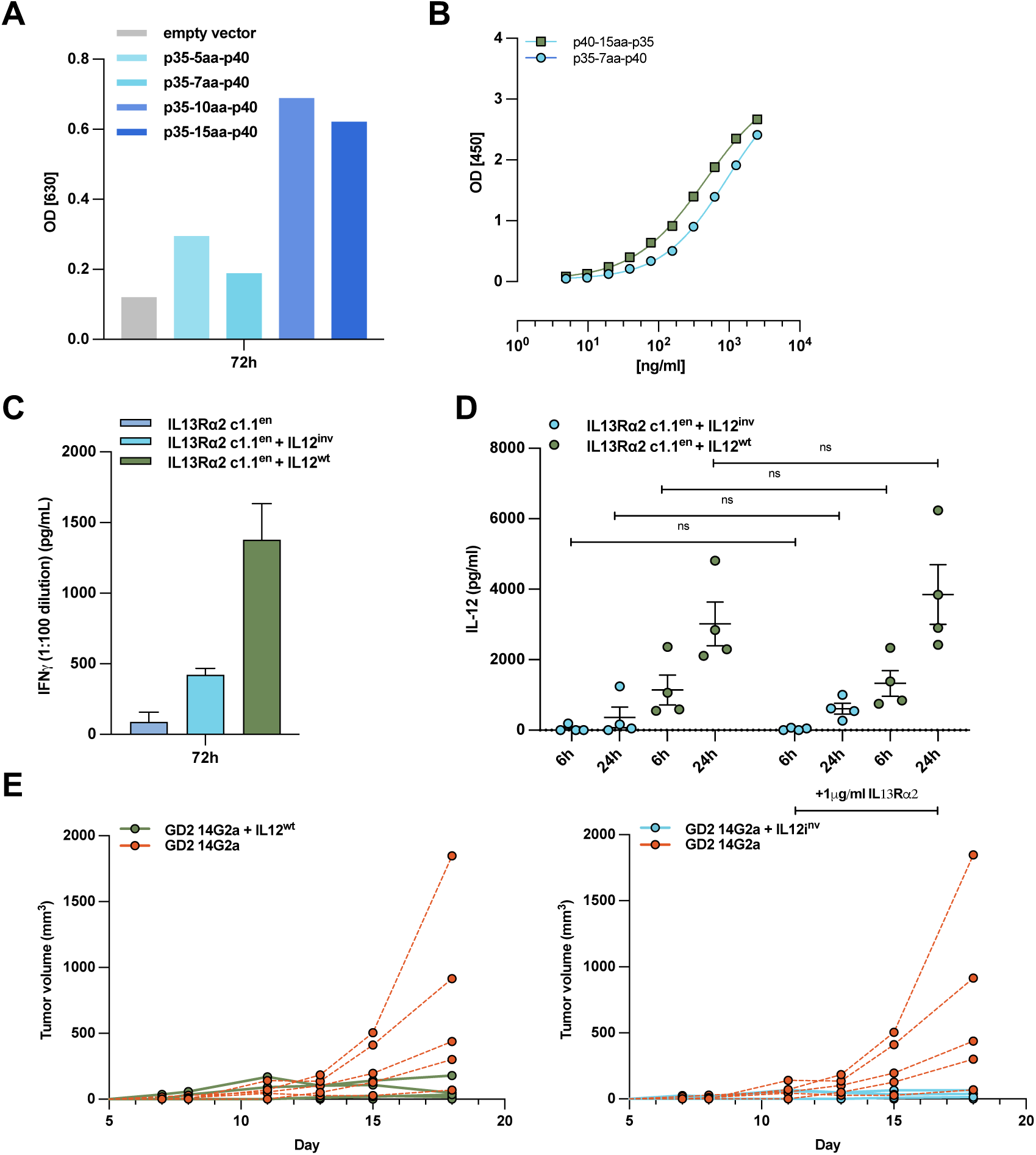
scIL12inv demonstrates enhanced safety while preserving its potent anti-cancer effects. **A.** Quantification of the SEAP secreted by HEK-Blue™ IL12 cultured in the presence of Jurkat cells expressing the p35 subunit followed by the p40 and connected with the indicated linker lengths for 72h. Results are expressed as representative of three independent experiments. **B.** Affinity of the designed p40-15aa-p35 and p35-7aa-p40 scIL12s to the immobilised IL12β2 was evaluated by ELISA assay (related to Figure 3c). **C.** ELISA detection of IFNγ production by NK cells co-cultured with Jurkat cells expressing IL13Rα2 c1.1en, IL13Rα2 c1.1en+IL12wt or IL13Rα2 c1.1en+IL12inv CAR for 72h (related to Figure **3G**). Cohorts are shown as mean ± SEM **D.** ELISA detection of IL12 production quantified by p70 analysis following IL13Rα2 stimulation of IL13Rα2 c1.1en+IL12wt or IL13Rα2 c1.1en+IL12inv CAR-T cells for the indicated time. **E.** Tumour volume measurement of each mouse used in the B16.F10-GD2 *in vivo* study (related to Figure **3J**). *P < 0.05, **P < 0.01, ***P < 0.001, ****P < 0.0001 and not significant (ns) P > 0.05.

**Supplementary Figure S4:**
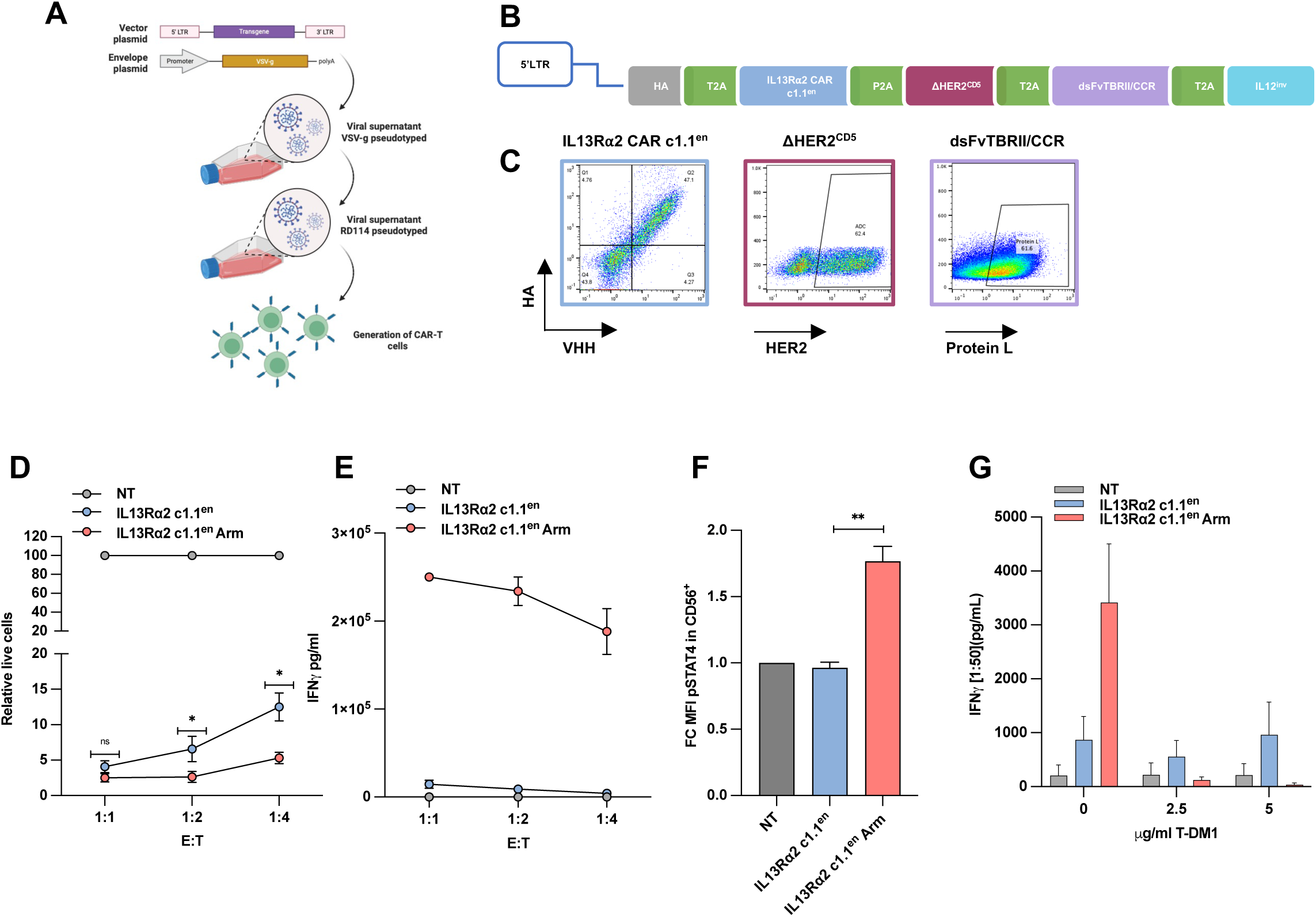
IL13Rα2 c1.1en Arm CAR shows resistance to TGF-β and enhanced cell persistence *in vitro* and *in vivo*. **A.** Schematic showing CAR vector with armouring modules. The vectors include an N-terminal transduction marker (HA) followed by the IL13Rα2 c1.1en. The additional armoured modules were inserted in the indicated order and separated by self-cleaving 2A peptides. LTR, long terminal repeat. **B.** Representative flow cytometry plots showing the detection of the IL13Rα2 c1.1en CAR and armoured modules. Results are expressed as representative of three independent donors for lymphocytes. **C.** Schematic representation of the transduction protocol created with Biorender.com. **D.** Viable target SNB19 cells following co-culture with IL13Rα2 c1.1en or IL13Rα2 c1.1en Arm CAR-T cells at the indicated E:T ratio quantified by flow cytometry. The percentage of live target cells was calculated by normalizing to the NT group. **E.** Quantification of IFN-γ production of CAR-T cells co-cultured with SNB19 target cells (related to Figure **5C**). **F.** pSTAT4 level in CD56+ cells following co-culture with CAR-T cells expressing IL13Rα2 c1.1en or IL13Rα2 c1.1en Arm for 72h. **G.** ELISA detection of IFNγ production of CAR-T cells co-cultured with SupT1-IL13Rα2 target cells following pre-exposure to T-DM1 at the indicated concentration (related to Figure **5F**).

**Supplementary Figure S5:**
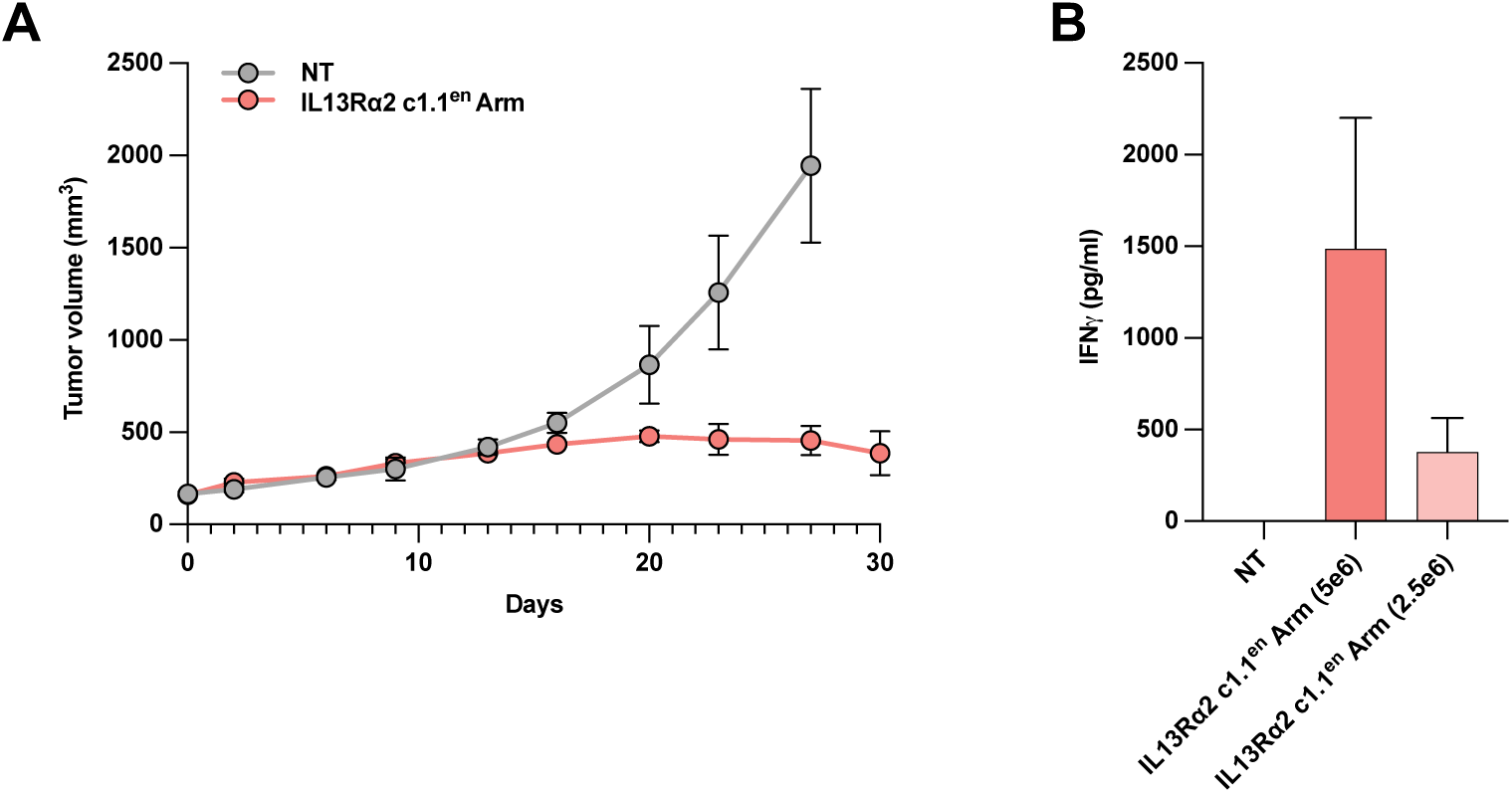
IL13Rα2 c1.1en Arm CAR shows enhanced cell persistence *in vivo*. **A.** Tumor volume measurements of NSG mice subcutaneously injected with U87-IL13Rα2 target cells, which were allowed to engraft for 8 days before a 2.5×106 dose of non-transduced T or CAR-T cells was intravenously injected via the tail vein. **B.** Quantification of IFN-γ levels in plasma samples collected from mice at day 30 post-treatment. IFN-γ concentrations were measured by ELISA.

## REFERENCES

1. Wu, W., Klockow, J.L., Zhang, M., Lafortune, F., Chang, E., Jin, L., Wu, Y., and Daldrup-Link, H.E. (2021). Glioblastoma multiforme (GBM): An overview of current therapies and mechanisms of resistance. Pharmacol. Res. 171, 105780. 10.1016/j.phrs.2021.105780.

2. Rodgers, L.T., Villano, J.L., Hartz, A.M.S., and Bauer, B. (2024). Glioblastoma Standard of Care: Effects on Tumor Evolution and Reverse Translation in Preclinical Models. Cancers 16, 2638. 10.3390/cancers16152638.

3. Land, C.A., Musich, P.R., Haydar, D., Krenciute, G., and Xie, Q. (2020). Chimeric antigen receptor T-cell therapy in glioblastoma: charging the T cells to fight. J. Transl. Med. 18, 428. 10.1186/s12967-020-02598-0.

4. Wykosky, J., Gibo, D.M., Stanton, C., and Debinski, W. (2008). Interleukin-13 receptor alpha 2, EphA2, and Fos-related antigen 1 as molecular denominators of high-grade astrocytomas and specific targets for combinatorial therapy. Clin. Cancer Res. Off. J. Am. Assoc. Cancer Res. 14, 199–208. 10.1158/1078-0432.CCR-07-1990.

5. Zeng, J., Zhang, J., Yang, Y.-Z., Wang, F., Jiang, H., Chen, H.-D., Wu, H.-Y., Sai, K., and Hu, W.-M. (2020). IL13RA2 is overexpressed in malignant gliomas and related to clinical outcome of patients. Am. J. Transl. Res. 12, 4702–4714.

6. Saikali, S., Avril, T., Collet, B., Hamlat, A., Bansard, J.-Y., Drenou, B., Guegan, Y., and Quillien, V. (2007). Expression of nine tumour antigens in a series of human glioblastoma multiforme: interest of EGFRvIII, IL-13Ralpha2, gp100 and TRP-2 for immunotherapy. J. Neurooncol. 81, 139–148. 10.1007/s11060-006-9220-3.

7. Brown, C.E., Warden, C.D., Starr, R., Deng, X., Badie, B., Yuan, Y.-C., Forman, S.J., and Barish, M.E. (2013). Glioma IL13Rα2 Is Associated with Mesenchymal Signature Gene Expression and Poor Patient Prognosis. PLoS ONE 8, e77769. 10.1371/journal.pone.0077769.

8. Brown, C.E., Badie, B., Barish, M.E., Weng, L., Ostberg, J.R., Chang, W.-C., Naranjo, A., Starr, R., Wagner, J., Wright, C., et al. (2015). Bioactivity and Safety of IL13Rα2-Redirected Chimeric Antigen Receptor CD8+ T Cells in Patients with Recurrent Glioblastoma. Clin. Cancer Res. 21, 4062–4072. 10.1158/1078-0432.CCR-15-0428.

9. Brown, C.E., Alizadeh, D., Starr, R., Weng, L., Wagner, J.R., Naranjo, A., Ostberg, J.R., Blanchard, M.S., Kilpatrick, J., Simpson, J., et al. (2016). Regression of Glioblastoma after Chimeric Antigen Receptor T-Cell Therapy. N. Engl. J. Med. 375, 2561–2569. 10.1056/NEJMoa1610497.

10. Bagley, S.J., Logun, M., Fraietta, J.A., Wang, X., Desai, A.S., Bagley, L.J., Nabavizadeh, A., Jarocha, D., Martins, R., Maloney, E., et al. (2024). Intrathecal bivalent CAR T cells targeting EGFR and IL13Rα2 in recurrent glioblastoma: phase 1 trial interim results. Nat. Med. 30, 1320–1329. 10.1038/s41591-024-02893-z.

11. Brown, C.E., Hibbard, J.C., Alizadeh, D., Blanchard, M.S., Natri, H.M., Wang, D., Ostberg, J.R., Aguilar, B., Wagner, J.R., Paul, J.A., et al. (2024). Locoregional delivery of IL-13Rα2-targeting CAR-T cells in recurrent high-grade glioma: a phase 1 trial. Nat. Med. 30, 1001–1012. 10.1038/s41591-024-02875-1.

12. Yang, L., Pang, Y., and Moses, H.L. (2010). TGF-beta and immune cells: an important regulatory axis in the tumor microenvironment and progression. Trends Immunol. 31, 220–227. 10.1016/j.it.2010.04.002.

13. Thomas, D.A., and Massagué, J. (2005). TGF-beta directly targets cytotoxic T cell functions during tumor evasion of immune surveillance. Cancer Cell 8, 369–380. 10.1016/j.ccr.2005.10.012.

14. Kloss, C.C., Lee, J., Zhang, A., Chen, F., Melenhorst, J.J., Lacey, S.F., Maus, M.V., Fraietta, J.A., Zhao, Y., and June, C.H. (2018). Dominant-Negative TGF-β Receptor Enhances PSMA-Targeted Human CAR T Cell Proliferation And Augments Prostate Cancer Eradication. Mol. Ther. J. Am. Soc. Gene Ther. 26, 1855–1866. 10.1016/j.ymthe.2018.05.003.

15. Car, B.D., Eng, V.M., Lipman, J.M., and Anderson, T.D. (1999). The Toxicology of Interleukin-12: A Review. Toxicol. Pathol. 27, 58–63. 10.1177/019262339902700112.

16. Jenks, S. (1996). After Initial Setback, IL-12 Regaining Popularity. JNCI J. Natl. Cancer Inst. 88, 576–577. 10.1093/jnci/88.9.576.

17. Jia, Z., Ragoonanan, D., Mahadeo, K.M., Gill, J., Gorlick, R., Shpal, E., and Li, S. (2022). IL12 immune therapy clinical trial review: Novel strategies for avoiding CRS-associated cytokines. Front. Immunol. 13, 952231. 10.3389/fimmu.2022.952231.

18. Asaadi, Y., Jouneghani, F.F., Janani, S., and Rahbarizadeh, F. (2021). A comprehensive comparison between camelid nanobodies and single chain variable fragments. Biomark. Res. 9, 87. 10.1186/s40364-021-00332-6.

19. Ramon, A., Ali, M., Atkinson, M., Saturnino, A., Didi, K., Visentin, C., Ricagno, S., Xu, X., Greenig, M., and Sormanni, P. (2024). Assessing antibody and nanobody nativeness for hit selection and humanization with AbNatiV. Nat. Mach. Intell. 6, 74–91. 10.1038/s42256-023-00778-3.

20. Marofi, F., Motavalli, R., Safonov, V.A., Thangavelu, L., Yumashev, A.V., Alexander, M., Shomali, N., Chartrand, M.S., Pathak, Y., Jarahian, M., et al. (2021). CAR T cells in solid tumors: challenges and opportunities. Stem Cell Res. Ther. 12, 81. 10.1186/s13287-020-02128-1.

21. Yang, L., Pang, Y., and Moses, H.L. (2010). TGF-beta and immune cells: an important regulatory axis in the tumor microenvironment and progression. Trends Immunol. 31, 220–227. 10.1016/j.it.2010.04.002.

22. Han, J., Alvarez-Breckenridge, C.A., Wang, Q.-E., and Yu, J. (2015). TGF-β signaling and its targeting for glioma treatment. Am. J. Cancer Res. 5, 945–955.

23. Golán-Cancela, I., and Caja, L. (2024). The TGF-β Family in Glioblastoma. Int. J. Mol. Sci. 25, 1067. 10.3390/ijms25021067.

24. Tugues, S., Burkhard, S.H., Ohs, I., Vrohlings, M., Nussbaum, K., Vom Berg, J., Kulig, P., and Becher, B. (2015). New insights into IL-12-mediated tumor suppression. Cell Death Differ. 22, 237–246. 10.1038/cdd.2014.134.

25. Lieschke, G.J., Rao, P.K., Gately, M.K., and Mulligan, R.C. (1997). Bioactive murine and human interleukin-12 fusion proteins which retain antitumor activity in vivo. Nat. Biotechnol. 15, 35–40. 10.1038/nbt0197-35.

26. Chen, X., Zaro, J.L., and Shen, W.-C. (2013). Fusion protein linkers: property, design and functionality. Adv. Drug Deliv. Rev. 65, 1357–1369. 10.1016/j.addr.2012.09.039.

27. Glassman, C.R., Mathiharan, Y.K., Jude, K.M., Su, L., Panova, O., Lupardus, P.J., Spangler, J.B., Ely, L.K., Thomas, C., Skiniotis, G., et al. (2021). Structural basis for IL-12 and IL-23 receptor sharing reveals a gateway for shaping actions on T versus NK cells. Cell 184, 983–999.e24. 10.1016/j.cell.2021.01.018.

28. Bloch, Y., Felix, J., Merceron, R., Provost, M., Symakani, R.A., De Backer, R., Lambert, E., Mehdipour, A.R., and Savvides, S.N. (2024). Structures of complete extracellular receptor assemblies mediated by IL-12 and IL-23. Nat. Struct. Mol. Biol. 31, 591–597. 10.1038/s41594-023-01190-6.

29. Lee, E.H.J., Murad, J.P., Christian, L., Gibson, J., Yamaguchi, Y., Cullen, C., Gumber, D., Park, A.K., Young, C., Monroy, I., et al. (2023). Antigen-dependent IL-12 signaling in CAR T cells promotes regional to systemic disease targeting. Nat. Commun. 14, 4737. 10.1038/s41467-023-40115-1.

30. Sahillioglu, A.C., and Schumacher, T.N. (2022). Safety switches for adoptive cell therapy. Curr. Opin. Immunol. 74, 190–198. 10.1016/j.coi.2021.07.002.

31. García-Alonso, S., Ocaña, A., and Pandiella, A. (2020). Trastuzumab Emtansine: Mechanisms of Action and Resistance, Clinical Progress, and Beyond. Trends Cancer 6, 130–146. 10.1016/j.trecan.2019.12.010.

32. Voisinne, G., Gonzalez de Peredo, A., and Roncagalli, R. (2018). CD5, an Undercover Regulator of TCR Signaling. Front. Immunol. 9, 2900. 10.3389/fimmu.2018.02900.

33. Bulcha, J.T., Wang, Y., Ma, H., Tai, P.W.L., and Gao, G. (2021). Viral vector platforms within the gene therapy landscape. Signal Transduct. Target. Ther. 6, 53. 10.1038/s41392-021-00487-6.

34. Sandrin, V., Boson, B., Salmon, P., Gay, W., Nègre, D., Le Grand, R., Trono, D., and Cosset, F.-L. (2002). Lentiviral vectors pseudotyped with a modified RD114 envelope glycoprotein show increased stability in sera and augmented transduction of primary lymphocytes and CD34+ cells derived from human and nonhuman primates. Blood 100, 823–832. 10.1182/blood-2001-11-0042.

35. Ghani, K., Cottin, S., De Campos-Lima, P.O., Caron, M., and Caruso, M. (2009). Characterization of an alternative packaging system derived from the cat RD114 retrovirus for gene delivery. J. Gene Med. 11, 664–669. 10.1002/jgm.1351.

36. Gumber, D., and Wang, L.D. (2022). Improving CAR-T immunotherapy: Overcoming the challenges of T cell exhaustion. eBioMedicine 77, 103941. 10.1016/j.ebiom.2022.103941.

37. Candolfi, M., Curtin, J.F., Nichols, W.S., Muhammad, A.G., King, G.D., Pluhar, G.E., McNiel, E.A., Ohlfest, J.R., Freese, A.B., Moore, P.F., et al. (2007). Intracranial glioblastoma models in preclinical neuro-oncology: neuropathological characterization and tumor progression. J. Neurooncol. 85, 133–148. 10.1007/s11060-007-9400-9.

38. Ludwig, J., and Hirschel, M. (2020). Methods and Process Optimization for Large-Scale CAR T Expansion Using the G-Rex Cell Culture Platform. In Chimeric Antigen Receptor T Cells Methods in Molecular Biology., K. Swiech, K. C. R. Malmegrim, and V. Picanço-Castro, eds. (Springer US), pp. 165–177. 10.1007/978-1-0716-0146-4_12.

39. Center for Biologics Evaluation and Research OTAT Town Hall: Gene Therapy Chemistry, Manufacturing, and Controls.

40. Sallusto, F., Geginat, J., and Lanzavecchia, A. (2004). Central Memory and Effector Memory T Cell Subsets: Function, Generation, and Maintenance. Annu. Rev. Immunol. 22, 745–763. 10.1146/annurev.immunol.22.012703.104702.

41. Kahlon, K.S., Brown, C., Cooper, L.J.N., Raubitschek, A., Forman, S.J., and Jensen, M.C. (2004). Specific Recognition and Killing of Glioblastoma Multiforme by Interleukin 13-Zetakine Redirected Cytolytic T Cells. Cancer Res. 64, 9160–9166. 10.1158/0008-5472.CAN-04-0454.

42. Krebs, S., Chow, K.K.H., Yi, Z., Rodriguez-Cruz, T., Hegde, M., Gerken, C., Ahmed, N., and Gottschalk, S. (2014). T cells redirected to interleukin-13Rα2 with interleukin-13 mutein–chimeric antigen receptors have anti-glioma activity but also recognize interleukin-13Rα1. Cytotherapy 16, 1121–1131. 10.1016/j.jcyt.2014.02.012.

43. Brown, C.E., Aguilar, B., Starr, R., Yang, X., Chang, W.-C., Weng, L., Chang, B., Sarkissian, A., Brito, A., Sanchez, J.F., et al. (2018). Optimization of IL13Rα2-Targeted Chimeric Antigen Receptor T Cells for Improved Anti-tumor Efficacy against Glioblastoma. Mol. Ther. 26, 31–44. 10.1016/j.ymthe.2017.10.002.

44. Krenciute, G., Krebs, S., Torres, D., Wu, M.-F., Liu, H., Dotti, G., Li, X.-N., Lesniak, M.S., Balyasnikova, I.V., and Gottschalk, S. (2016). Characterization and Functional Analysis of scFv-based Chimeric Antigen Receptors to Redirect T Cells to IL13Rα2-positive Glioma. Mol. Ther. 24, 354–363. 10.1038/mt.2015.199.

45. Choi, B.D., Gerstner, E.R., Frigault, M.J., Leick, M.B., Mount, C.W., Balaj, L., Nikiforow, S., Carter, B.S., Curry, W.T., Gallagher, K., et al. (2024). Intraventricular CARv3-TEAM-E T Cells in Recurrent Glioblastoma. N. Engl. J. Med. 390, 1290–1298. 10.1056/NEJMoa2314390.

46. Turley, S.J., Cremasco, V., and Astarita, J.L. (2015). Immunological hallmarks of stromal cells in the tumour microenvironment. Nat. Rev. Immunol. 15, 669–682. 10.1038/nri3902.

47. Tang, N., Cheng, C., Zhang, X., Qiao, M., Li, N., Mu, W., Wei, X.-F., Han, W., and Wang, H. (2020). TGF-β inhibition via CRISPR promotes the long-term efficacy of CAR T cells against solid tumors. JCI Insight 5, e133977. 10.1172/jci.insight.133977.

48. DeRenzo, C., and Gottschalk, S. (2019). Genetic Modification Strategies to Enhance CAR T Cell Persistence for Patients With Solid Tumors. Front. Immunol. 10, 218. 10.3389/fimmu.2019.00218.

49. Shum, T., Omer, B., Tashiro, H., Kruse, R.L., Wagner, D.L., Parikh, K., Yi, Z., Sauer, T., Liu, D., Parihar, R., et al. (2017). Constitutive Signaling from an Engineered IL7 Receptor Promotes Durable Tumor Elimination by Tumor-Redirected T Cells. Cancer Discov. 7, 1238–1247. 10.1158/2159-8290.CD-17-0538.

50. Lin, F.Y., Stuckert, A., Tat, C., White, M., Ruggieri, L., Zhang, H., Mehta, B., Lapteva, N., Mei, Z., Major, A., et al. (2024). Phase I Trial of GD2.CART Cells Augmented With Constitutive Interleukin-7 Receptor for Treatment of High-Grade Pediatric CNS Tumors. J. Clin. Oncol. 42, 2769–2779. 10.1200/JCO.23.02019.

51. Seder, R.A., Gazzinelli, R., Sher, A., and Paul, W.E. (1993). Interleukin 12 acts directly on CD4+ T cells to enhance priming for interferon gamma production and diminishes interleukin 4 inhibition of such priming. Proc. Natl. Acad. Sci. 90, 10188–10192. 10.1073/pnas.90.21.10188.

52. Gazzinelli, R.T., Wysocka, M., Hayashi, S., Denkers, E.Y., Hieny, S., Caspar, P., Trinchieri, G., and Sher, A. (1994). Parasite-induced IL-12 stimulates early IFN-gamma synthesis and resistance during acute infection with Toxoplasma gondii. J. Immunol. Baltim. Md 1950 153, 2533–2543.

53. Lasek, W., Zagożdżon, R., and Jakobisiak, M. (2014). Interleukin 12: still a promising candidate for tumor immunotherapy? Cancer Immunol. Immunother. CII 63, 419–435. 10.1007/s00262-014-1523-1.

54. Zhang, L., Morgan, R.A., Beane, J.D., Zheng, Z., Dudley, M.E., Kassim, S.H., Nahvi, A.V., Ngo, L.T., Sherry, R.M., Phan, G.Q., et al. (2015). Tumor-Infiltrating Lymphocytes Genetically Engineered with an Inducible Gene Encoding Interleukin-12 for the Immunotherapy of Metastatic Melanoma. Clin. Cancer Res. 21, 2278–2288. 10.1158/1078-0432.CCR-14-2085.

55. Zhang, L., Davies, J.S., Serna, C., Yu, Z., Restifo, N.P., Rosenberg, S.A., Morgan, R.A., and Hinrichs, C.S. (2020). Enhanced efficacy and limited systemic cytokine exposure with membrane-anchored interleukin-12 T-cell therapy in murine tumor models. J. Immunother. Cancer 8, e000210. 10.1136/jitc-2019-000210.

56. Hu, J., Yang, Q., Zhang, W., Du, H., Chen, Y., Zhao, Q., Dao, L., Xia, X., Natalie Wall, F., Zhang, Z., et al. (2022). Cell membrane-anchored and tumor-targeted IL-12 (attIL12)-T cell therapy for eliminating large and heterogeneous solid tumors. J. Immunother. Cancer 10, e003633. 10.1136/jitc-2021-003633.

57. Lee, E.H.J., Murad, J.P., Christian, L., Gibson, J., Yamaguchi, Y., Cullen, C., Gumber, D., Park, A.K., Young, C., Monroy, I., et al. (2023). Antigen-dependent IL-12 signaling in CAR T cells promotes regional to systemic disease targeting. Nat. Commun. 14, 4737. 10.1038/s41467-023-40115-1.

58. Zhou, X., Dotti, G., Krance, R.A., Martinez, C.A., Naik, S., Kamble, R.T., Durett, A.G., Dakhova, O., Savoldo, B., Di Stasi, A., et al. (2015). Inducible caspase-9 suicide gene controls adverse effects from alloreplete T cells after haploidentical stem cell transplantation. Blood 125, 4103–4113. 10.1182/blood-2015-02-628354.

59. Del Bufalo, F., De Angelis, B., Caruana, I., Del Baldo, G., De Ioris, M.A., Serra, A., Mastronuzzi, A., Cefalo, M.G., Pagliara, D., Amicucci, M., et al. (2023). GD2-CART01 for Relapsed or Refractory High-Risk Neuroblastoma. N. Engl. J. Med. 388, 1284–1295. 10.1056/NEJMoa2210859.

60. Diaconu, I., Ballard, B., Zhang, M., Chen, Y., West, J., Dotti, G., and Savoldo, B. (2017). Inducible Caspase-9 Selectively Modulates the Toxicities of CD19-Specific Chimeric Antigen Receptor-Modified T Cells. Mol. Ther. J. Am. Soc. Gene Ther. 25, 580–592. 10.1016/j.ymthe.2017.01.011.

